# Impact of genomic background and developmental state on signaling pathways and response to therapy in glioblastoma patient-derived cells

**DOI:** 10.1101/2024.03.14.585115

**Authors:** Artem Berezovsky, Oluwademilade Nuga, Indrani Datta, Kimberly Bergman, Thais Sabedot, Katherine Gurdziel, Susan Irtenkauf, Laura Hasselbach, Yuling Meng, Claudius Mueller, Emanuel F. Petricoin, Stephen Brown, Neeraja Purandare, Sidhesh Aras, Tom Mikkelsen, Laila Poisson, Houtan Noushmehr, Douglas Ruden, Ana C. deCarvalho

## Abstract

Glioblastoma (GBM) tumors represents diverse genomic epigenomic, and transcriptional landscapes, with significant intratumoral heterogeneity that challenges standard of care treatments involving radiation (RT) and the DNA-alkylating agent temozolomide (TMZ). In this study, we employed targeted proteomics to assess the response of a genomically-diverse panel of GBM patient-derived cancer stem cells (CSCs) to astrocytic differentiation, growth factor withdrawal and traditional high fetal bovine serum culture. Our findings revealed a complex crosstalk and co-activation of key oncogenic signaling in CSCs and diverse patterns of response to these external stimuli. Using RNA sequencing and DNA methylation, we observed common adaptations in response to astrocytic differentiation of CSCs across genomically distinct models, including BMP-Smad pathway activation, reduced cholesterol biosynthesis, and upregulation of extracellular matrix components. Notably, we observed that these differentiated CSC progenies retained a subset of stemness genes and the activation of cell survival pathways. We also examined the impact of differentiation state and genomic background on GBM cell sensitivity and transcriptional response to TMZ and RT. Differentiation of CSCs increased resistance to TMZ but not to RT. While transcriptional responses to these treatments were predominantly regulated by p53 in wild-type p53 GBM cells, its transcriptional activity was modulated by the differentiation status and treatment modality. Both mutant and wild-type p53 models exhibited significant activation of a DNA-damage associated interferon response in CSCs and differentiated cells, suggesting this pathway may play a wider role in GBM response to TMZ and RT. Our integrative analysis of the impact of GBM cell developmental states, in the context of genomic and molecular diversity of patient-derived models, provides valuable insights for pre-clinical studies aimed at optimizing treatment strategies.

## Introduction

Glioblastoma (GBM), classified as World Health Organization (WHO) Grade 4 isocitrate dehydrogenase (IDH)-wildtype astrocytoma, remains the most aggressive and prevalent primary brain tumor in adults [1]. Since 2005, standard of care treatment of GBM includes maximal safe surgical resection [2], followed by ionizing radiation therapy (RT) and the brain-penetrant DNA-alkylating agent temozolomide (TMZ) [3]. Surgery alone is not curative [2] and TMZ provides a modest, significant increase in median overall survival in patients with newly diagnosed GBMs when combined with RT compared to RT alone [4]. TMZ sensitivity in newly diagnosed GBMs is modulated by the expression of DNA repair protein, O-6-methylguanine-DNA methyltransferase (MGMT). The silencing of MGMT through promoter hypermethylation is a predictive marker of TMZ sensitivity for newly diagnosed GBMs [5], while MGMT expression is also regulated by DNA methylation independent mechanisms [6, 7]. Most GBM patients have known actionable therapeutical targets [8], clinical trials for targeted therapies have not demonstrated sufficient efficacy in improving overall and progression-free survival [9].

Intra-tumoral heterogeneity (ITH) is a key factor contributing to GBM resistance to treatment and high recurrence rate. Genomically-defined subclonal populations presenting differential sensitivity to pharmacological agents can arise in post-treatment recurrent GBMs [10]. Another contributor to ITH is oncogene amplification in extrachromosomal DNA (ecDNA), as we previously reported cases of low frequency ecDNA in newly diagnosed GBM exhibiting a notable increase in prevalence at recurrence following treatment [11]. Non-genetic factors also contribute to ITH, such as the high degree of plasticity of GBM cells, reflected in the ability to transition into a continuum of cell states analogous to neural development, from neural stem/progenitor-like to differentiated astrocyte-like cells, as shown by bulk tumor RNA sequencing (seq) deconvolution and single cell RNAseq analyses [12]. ITH involves dynamic shifts in subclonal composition in response to treatment and to changes in the microenvironment in GBM [13, 14]. Thus, it is important to consider ITH when designing preclinical studies to test optimization of treatment efficacy for GBMs.

Tumor cell subpopulations exhibiting cancer stem cell-like (CSC) properties contribute to another layer of ITH through developmental cell state plasticity [15]. Long-term self-renewal in culture and the ability to differentiate and recapitulate the original tumor upon orthotopic implantation in immunocompromised rodents are defining attributes of cancer stem cells (CSCs) [16, 17]. These properties make CSCs invaluable for developing GBM patient-derived models for pre-clinical studies. An established strategy to select for GBM CSC from surgical samples involves culturing dissociated tumor cells in selective serum-free media, originally formulated for the isolation of mouse neural progenitor cells, resulting in “neurosphere” cultures [18, 19]. In recent years, GBM neurosphere cultures have been extensively validated as a renewable source of patient-derived neoplastic cells retaining remarkable genomic fidelity to the original tumor [11, 20, 21]. These neurosphere/CSC cultures are amenable to in vitro biological and experimental therapeutics studies [22] and are suitable for validation in xenografts. To enhance the translational applicability of in vitro investigations aimed at optimizing treatment efficacy in GBMs, it is imperative to consider the influence of patient genomic diversity and intra-tumoral phenotypic variability. Ultimately, the diversity in genomic and epigenomic drivers among GBM patients, along with the additional variability associated with dynamic ITH, converge on the deregulation of key oncogenic signaling involving receptor tyrosine kinases (RTK), phosphoinositide 3-kinases (PI3K)- protein kinase B (AKT)- mammalian target of rapamycin (mTOR) and mitogen-activated protein kinase (MAPK) pathways. These pathways are deregulated in 90% of GBM patients [23], and cross activation of PI3K and MAPK contribute to several aspects of GBM malignancy [24].

The objective of this study is to investigate the contributions of somatic genomic alterations in conjunction with GBM cellular developmental states to the modulation of key signaling pathways and response to TMZ and RT. Employing targeted proteomics, we assessed changes in the steady-state levels of key oncogenic signaling components, following a 2-week exposure of CSCs to astrocytic differentiation (SDCs), growth factor withdrawal, or traditional culture media supplemented with 10% FBS. in a genomically-diverse panel of CSCs derived from newly diagnosed GBM patients [11, 25, 26]. The representation of various genomic backgrounds and external stimuli in the targeted proteomics dataset provided insights into the complex signaling networks co-activated in GBMs, while uncovering a strong positive correlation between NF-kB activation and MGMT protein expression, regardless of MGMT promoter methylation status. Some of the signaling and transcriptional program adaptations in response to astrocytic differentiation of CSCs were shared among the genomically distinct models, including BMP-Smad pathway activation, decrease in cholesterol biosynthesis, increase in extracellular matrix (ECM) production, and retention of the expression of a subset of stemness genes. Other CSC differentiation mediated alterations were tumor specific, such as activation of the interferon (IFN) response. Here, we show that differentiation of CSCs increases resistance to TMZ but not to RT, and that transcriptional responses to these treatments are largely regulated by p53 in wildtype (wt) p53 GBM cells and modulated by cell differentiation state. In both mutant and wt p53 models, we observed prominent activation of DNA-damage associated IFN response. We show that modeling the differentiation of GBM CSCs in vitro provides a powerful platform to uncover the interaction of genomic landscape and cell state on signaling pathways and response to treatment.

## Results

### Alterations in key cell signaling pathways in glioblastoma cancer stem cells in response to astrocytic differentiation, growth factor withdrawal and traditional 10% FBS media

CSCs derived from 8 newly diagnosed GBM patients, representing the main genomic drivers were selected for this study (Fig 1A). These CSC lines meet the requirement of the two defining stemness criteria: long term self-renewal and orthotopic tumor formation in immunocompromised mice [11, 25, 26]. FBS, a supplement supporting the culture of a variety of mammalian cells, contains hormones, growth factors, cell attachment factors and other nutrients [27]. Media supplemented with 10% FBS had been widely employed to propagate glioma cells in culture, prior to seminal data demonstrating genetic drift occurring from serial passaging under these conditions [28]. In contrast, 1-2% FBS added to defined media has been successfully used to differentiate human-induced pluripotent stem cells (hiPSC)-derived neural stem cells into astrocytic lineage [29, 30]. Using this paradigm to model a differentiated phenotype in culture, CSCs growing in 3D cultures in defined neurosphere media, supplemented with, 20 ng/ml each epidermal (EGF) and basic fibroblast (bFGF) growth factors (NMGF), were differentiated into an astrocytic phenotype in NMGF supplemented with 2% FBS for 14 days. This differentiated CSC progeny is referred to here as “serum differentiated cells” (SDC). For comparison, we tested two additional growth conditions: growth factor-depleted neurosphere media (NM), or media supplemented with 10% FBS (10% FBS) (Fig 1B). After 2 weeks in culture, cell lysates were obtained and analyzed by Reverse Phase Protein Arrays (RPPA) in triplicates, as described [31, 32]. The levels of 66 proteins or post-translational modifications (PTM) in the cell lysates were quantified (S1 Table). Pairwise comparisons between NMGF (CSC) culture and either NM or 2% FBS (SDC) for the 8 cell lines did not show a global shift in the levels of protein/PTMs levels (S1_Fig), but important patterns of alteration in specific signaling were observed. Phosphorylation (phospho) levels of serine (Ser)-residues in receptor-dependent r-Smad1/5/8, which are primarily activated by BMP ligand signaling, were notably upregulated in SDCs in relation to CSCs in all 8 GBM models (Fig 2A). Consistent with bone morphogenetic protein (BMP) ligands being present in FBS [33], increased levels of p-Smad1/5/8 were also observed in 10% FBS cultures for all models (S1 Table). The tumor suppressor phosphatase and tensin homolog (PTEN) function is frequently lost in GBMs through gene deletion and mutations, as seen in 7/8 GBM tumors represented in this study (Fig 1A), leading to activation of AKT[34]. AKT phosphorylation at both Y308 (by pyruvate dehydrogenase kinase 1 (PDK1)) and S473 (by mammalian target of rapamycin complex 2 (mTORC2)) was suppressed in the absence of growth factors (NM) for all lines, except for HF2927, which carries ligand independent epidermal growth factor receptor (EGFR) variant III (viii) amplification. In 2% FBS (SDCs), AKT phosphorylation was suppressed for HF2587, HF3035 and HF3077, and upregulated or unchanged in the remaining models (Fig 2A). In response to growth factor withdrawal, phosphorylation levels of S6 ribosomal proteins (RBS6), downstream of mTORC1, were decreased while phospho-extracellular signal-regulated kinases (ERK) increased for most CSC lines (Fig 2A). Phospho-EGFR (Y1173 and Y1068) levels were increased in response to growth factor withdrawal exclusively in the EGFRvIII line HF2927 (Fig 2A). Expression of MET proto-oncogene, receptor tyrosine kinase (cMET) was observed for all CSCs (S1 Table), but in the absence of MET ligand (hepatocyte growth factor) in the media, only HF2927 and HF3016 presented modest levels of MET activation, which was suppressed in 2% FBS (Fig 2A).

**Fig 1.**
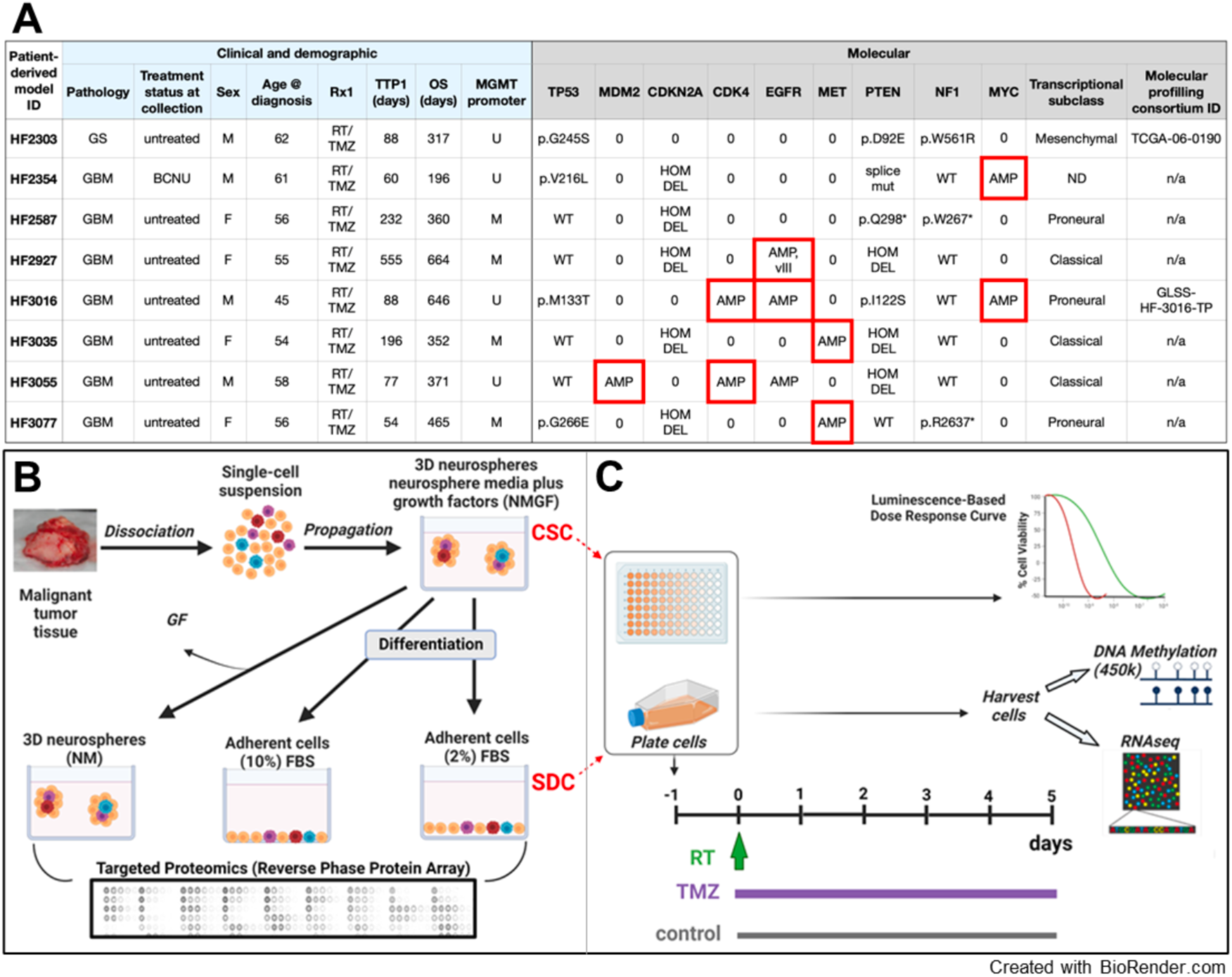
Glioblastoma patient-derived models and experimental design. **A)** Clinical and molecular data associated with the GBM patient-derived models. GS, Gliosarcoma; M/F, male/female; RT, radiation therapy; TMZ, temozolomide; Rx1, first-line therapy; TTP, time to progression; OS, overall survival; U/M, unmethylated/methylated MGMT promoter. Red outline denotes extrachromosomal (ecDNA) amplification. MDM2, Mouse double minute 2 homolog; CDKN2A, cyclin-dependent kinase inhibitor 2A; NF1, Neurofibromin 1; MYC family, MYC proto-oncogene, bHLH transcription factor; ND, not determined; AMP, amplification; mut, mutation; hom del, homozygous deletion; WT, wild type; 0, diploid. **B)** Schematic depicting neurosphere culture in NMGF media for selection and amplification of cancer stem cells (CSCs) from surgical specimens, followed by 2 weeks incubation in altered conditions: withdrawal of growth factors (NM), and addition of 2% or 10% FBS. For the purposes of this study, serum-differentiated cells (SDC) are CSCs progeny cultured in NMGF supplemented with 2% FBS. The effect of the different culture conditions on cell signaling were compared by RPPA. **C)** The sensitivity of CSC and SDC to single dose radiation (RT) or 5-day TMZ was measured in three select models. Transcriptional and epigenomic reprograming in SDC vs CSC, and in response to treatment were evaluated by bulk RNAseq and 450k DNA methylation array.

**Fig 2.**
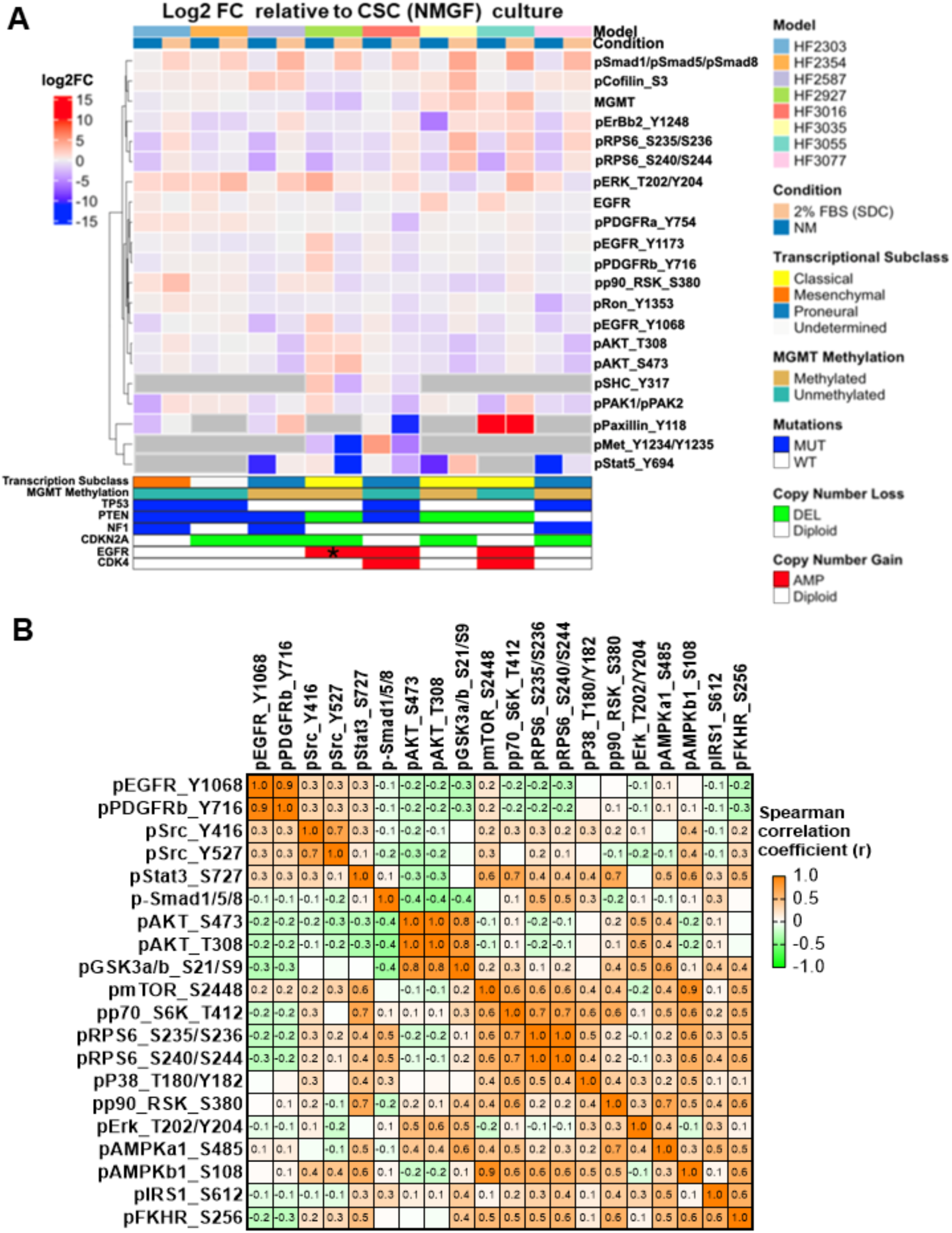
Targeted proteomics reveals common and cell specific alterations in key signaling CSCs in response to differentiation and identifies interdependencies in activation of key signaling mediators. **A)** Heatmap depicting log2 fold change (FC) relative to NMGF (CSC) for the top 5% most variable proteins/PTM. Each value represents mean log2FC for n=3 RPPA measurements/group. Proteins/PTMs which were not detected under all three culture conditions were filtered out for each model (grey cells). MUT, mutation; WT, wild type; AMP, amplification; DEL, deletion **B)** Heatmap of Spearman correlation coefficient values comparing phosphorylation levels in key cell signaling mediators in 8 patient derived GBM models grown in 4 media conditions in triplicates.

We used this unique targeted proteomics dataset, integrating various genomic backgrounds and culture conditions, to further investigate the signaling networks in glioblastoma. The Spearman correlation of key proteins and post-translational modifications levels in all 96 samples (8 GBM lines in 4 media conditions in triplicate) was calculated and displayed in the heatmap (Fig 2B). Glycogen synthase kinase-3 (GSK3)α/β is a multifunctional kinase frequently inhibited by phosphorylation downstream of PI3K/AKT, mTORC1 or MAPK signaling. Here, we observed that phosphorylation of GSK3α/β at S21/S9 was strongly correlated with AKT activation (Fig 2B). While mTORC1 is activated by AKT [35], in this dataset AKT activation did not correlate with markers of mTORC1 signaling, such as mTOR phosphorylation at the C-terminus S2448 residue [36], p70 ribosomal protein S6 kinase 1 (S6K) phosphorylation at T412, or phosphorylation of RBS6 [37] (Fig 2B). On the other hand, phosphorylation of AMP-activated protein kinase (AMPK) subunit β1 at S108 was highly correlated with mTORC1 activation (Fig 2B). AMPK αβɣ heterotrimeric complex is activated in response to low oxygen, low nutrients, and DNA damage, playing important roles in cancer [37]. AMPK is inactivated by AKT phosphorylation of AMPKα1 at S485 residue, while phosphorylation at AMPKβ1 S108 sensitizes AMPK to agonists [38], having a function in pro-survival pathways [39]. SRC proto-oncogene, non-receptor tyrosine kinase (Src) is a node of convergence for receptor mediated signaling pathways, activating substrates in the MAPK, JAK/Stat3, and PI3K/AKT pathways to promote tumor cell survival, proliferation and invasion in various cancers, including GBMs [40]. Src activity is inhibited by phosphorylation of Y527 by C-terminal Src kinase (CSK) and regulated by dephosphorylation by protein tyrosine phosphatases (PTPs), leading to conformational change and autophosphorylation of Y416 [41]. Levels of inhibitory Src phospho-Y527 were elevated in HF2354, HF2587, HF3016 and HF3077 relative to other CSC lines while the levels of Src phospho-Y416 were less variable among CSC lines and media conditions (S1 Table) and positively correlated with Src phospho-Y527 (r=0.705, CI: 0.584 to 0.796) but not with phospho-signal transducer and activator of transcription 3 (Stat3 S727), a canonical downstream signal transducer (Spearman correlation, r=0.302, CI: 0.1026 to 0.4788) (Fig 2B). This bi-phosphorylation of Src has been previously reported to reflect an intermediate state between open and closed kinase conformations with blockage of the SH3 domain leading to limited Stat3 phosphorylation activity, suggesting that additional stimulatory factors are required for downstream signaling [42]. Phospho-Stat3 was positively correlated with mTORK1 (phospho-mTOR, phospho-SK6) and MAPK (phospho-RSK) signaling (Fig 2B). Signaling through insulin like growth factor 1 receptor (IGF1R) and insulin receptor (IR) promotes cancer cell proliferation, survival, and treatment resistance in diverse malignancies. Upon ligand binding, IGF1R homodimer or IGF1R/IR heterodimer receptors recruit insulin receptor substrate (IRS) adaptor protein to transduce downstream signaling through binding of src-homology-2 (SH2) proteins, such as p85 to activate PI3K and growth factor receptor bound protein 2 (GRB2) to activate ERK [43]. All four media conditions tested here contain insulin at a final concentration of 50 µg/ml (N2 supplement), which has been shown to activate IGF1R, although at a lower rate than the canonical ligand IGF2 [44], while the FBS supplemented media contains additional insulin and IGF1/2 ligands [27]. Consistently, activation of IGF1R (phospho-Y1135/Y1136) /IR (phospho-Y1150/Y1151) presented little variation among GBM lines and culture conditions (S1 Table). mTORC1 phosphorylates IRS1 at S616 residue leading to the adaptor turnover [45], but p-IRS1 did not correlate with mTORC1 activation. Instead, a strong correlation of p-IRS1 with inactivating phosphorylation of forkhead box protein O1 (FOXO1/FKHR) on S256, which is attributed to AKT [46], was observed (Fig 2B). This targeted proteomics analysis using genomically diverse patient samples in different culture conditions reveled a complex pattern in the integration of signaling cascade nodes, likely reflecting deregulation of multiple oncogenic pathways observed in GBMs.

Z-scores were calculated for each protein/PTM based on its distribution across all 8 models and 4 media conditions, and cumulative z-scores were determined for key signaling pathways and cellular functions (S1 Table). Interestingly, we observed a remarkable pattern of inhibition for key signaling pathway and functions in 10% FBS, in contrast with 2% FBS (SDC) cultures (Fig 3A), further corroborating the inadequacy of culturing GBM cells in 10% FBS [28, 47]. Receptor tyrosine kinase activation (“RTK”) index, based on the levels of Tyr phosphorylation corresponding to activation of 11 RTKs (S1 Table), decreased in 2% FBS (SDCs) relative to CSCs in HF2927 and HF3016, the only 2 models harboring EGFR amplification on extrachromosomal DNA (ecDNA) [11], and increased in the remaining 6 GBM models (Fig 3A). The levels of EGFR were 10-fold higher for HF2927 CSCs, which also carries the EGFRvIII variant (Fig 1A), relative to HF3016, while the levels of phospho-EGFR at carboxy-terminal Y1068 and Y1173 residues were comparable for these 2 CSC lines (Fig 3A). Interestingly, EGF withdrawal for 2 weeks (NM) led to a 4-fold increase in the levels of EGFR activation in HF2927 (S1 Table, Fig 2A), suggesting an inhibitory effect of long-term exposure to EGF ligand on EGFRvIII. RTK activation correlated with downstream RTK signaling pathway (“TK_Signaling”), which includes both MAPK and PI3K/Akt/mTOR (“PI3K”) pathways (Fig 3A). As part of RTK signaling convergence, SHC Adaptor Protein 1 (SHC1) binds to phosphorylated EGFR Y1173, which is further phosphorylated on Y239 and Y317 resulting in recruitment of Grb2 and downstream activation of MAPK and PI3K/AKT pathways [46]. Phospho-Shc1 (Y317) was only observed for HF2927 and HF3016 CSCs, and in both cases upregulated upon growth factor withdrawal (NM) and downregulated in the presence of FBS, following the pattern of EGFR activation (S1 Table, Fig 2A). MAPK signaling activation and cytoskeletal function, consisting of regulation of actin and microtube dynamics (S1 Table), were maintained or increased in SDCs for all models except HF2927 and HF3016 (Fig 3A), analogous to what we observed for the RTK index. The PI3K/AKT/mTOR pathway activation score was low for HF2927 CSCs, relative to the other CSC lines, and remained low in NM and 2% FBS, despite genomic loss of PTEN, EGFRvIII amplification, and high level of EGFR activation (Fig. 3A). PI3K/AKT/mTOR activation index was higher for the other 7 CSCs, maintained in SDCs, and decreased in the absence of growth factors (Fig. 3A).

**Fig 3.**
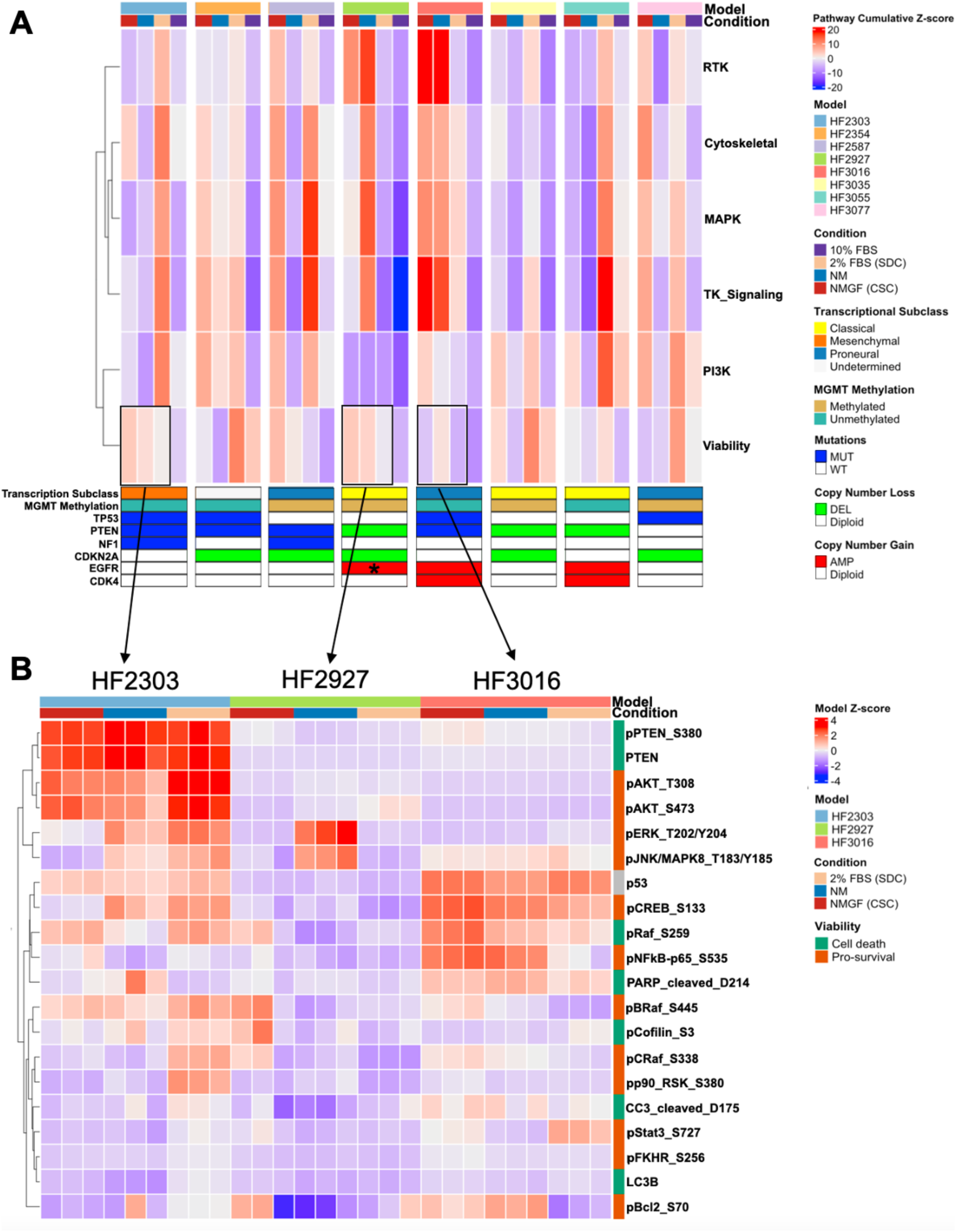
Comparative targeted proteomics of CSCs growing in 4 different conditions. **A)** Heatmap depicting mean signaling pathway activation for triplicate samples in 4 culture conditions, for CSCs derived from 8 patients with diverse genomic landscape, transcriptional subclass and MGMT promoter methylation. Hierarchical clustering of rows was based on Spearman correlation analysis. **B)** Heatmap for protein/PTM z-scores involved in cell death and survival signaling for 3 models grown in NMGF (CSC), NM and 2% FBS (SDC), in triplicate (boxes in (A)).

Since several of these signaling pathways are involved in cell survival and proliferation, we established a cell “viability index” measuring the net result from the sum of the level of protein/PTM associated with cell survival and proliferation minus the level of markers associated with programed cell death (S1 Table). We observed that the viability index varied among CSC lines and was not impacted by culture conditions in the same pattern, decreasing in SDC in half of the models and increasing in the other half. (Fig 3A). Three models, HF2303, HF2927 and HF3016, representing each of the 3 transcriptional subclasses, diverse genomic features (Fig 1A), and a range of viability index values, were selected for further characterization. Levels of individual protein/PTM associated with cell death and survival are shown (Fig 3B). High PTEN expression was observed in HF2303, harboring pathogenic p.D92E mutation in the WPD loop [48], corresponding to a high level of phosphorylation at S380 in the carboxy terminal tail, a crucial post-translational modification for blocking proteasomal degradation [49]. High levels of AKT activation observed for HF2303 CSCs were further increased in SDCs, indicating that the p.D92E mutation impairs PTEN ability to inhibit AKT. Conversely, PTEN levels were very low in HF3016, which carries a pathogenic mutation in the P-loop, p.I122S [48], comparable to PTEN-null HF2927 (Fig 3B). Increased levels of phospho-ERK and phospho- c-Jun N-terminal kinase (JNK) were observed for HF2303 and HF2927 in response to growth factor withdrawal (NM), and in HF2303, the increase was also observed in SDCs, relative to CSCs. The levels of p53 protein were consistent with the mutation status for these lines (Fig 3B). Thus, our comparative analysis confirms that at the individual protein/PTM levels, differences in pathways regulating cell survival and death in untreated GBM cells are largely determined by the genomic landscape, with cell state playing a lesser role.

### CSC presents increased sensitivity to temozolomide but not to radiation compared to their SDC progeny

MGMT epigenetic silencing is a clinically used biomarker to predict response to TMZ treatment. HF3016 and HF2303 CSCs were derived from tumors with unmethylated and HF2927 CSC from a tumor with methylated MGMT promoter (Fig. 1A). Accordingly, HF2927 cells presented the lowest level of MGMT protein. Despite both having unmethylated promoter status, MGMT levels were 2.5-fold higher for HF3016 relative to HF2303 (Fig. 4A, S1 Table). MGMT protein level in HF2303 was unchanged among all three culture conditions, while MGMT was significantly downregulated in SDCs relative to CSCs for HF2927 and HF3016 (Fig 4A). We confirmed that MGMT promoter was unmethylated for HF2303 and methylated for HF2927 cells and did not observe changes in methylation between CSC and SDCs for either of these models (S2 Fig). DNA methylation data is unavailable for HF3016 SDCs, preventing us from entirely ruling out differentiation-mediated alterations in MGMT promoter methylation in this model. We observed that MGMT protein expression pattern in the two models with unmethylated promoter, HF3016 and HF2303 (Fig 4A), was highly correlated (Spearman r =0.823; 95% CI: 0.6213 to 0.9228) with the levels of activated nuclear factor-kappa B (NF-kB) p65 (phospho-S536) (Fig 3B, S1 Table). Remarkably, the correlation between activated NF-kB and MGMT levels was significant for all models, regardless of promoter methylation status, but stronger for those presenting unmethylated MGMT (Fig 4B). To investigate to what extent the developmental state affected the sensitivity of GBM cells to genotoxic treatments, HF2303, HF2927 and HF3016 CSCs, along with their SDC progenies, were treated with TMZ and RT (Fig 1C). These models represent diverse molecular characteristics known to modulate response to DNA-damaging agents, such as TP53 status and MGMT promoter methylation. First, we compared cell proliferation rates between 3D CSC and 2D SDC cultures. SDCs proliferated at a higher rate than CSCs for HF3016 only, while no significant difference was observed for the other two lines (Fig 5A). Cells were then treated for 4 days with doses ranging from 0 (DMSO) to 400 mM TMZ, in quintuplicates. To compensate for possible binding of TMZ by proteins in FBS, even at the low concentration used, the serum-free neurosphere media was supplemented with bovine serum albumin (1%) [50]. Half maximal inhibitory concentration (IC50) and area above the curve (AAC) were calculated from dose-response curves generated by non-linear fitting. HF2927 cells were more sensitive to TMZ in comparison with the other two cell lines (Fig 5B), as expected based on MGMT promoter methylation and expression (Fig 4A). Differentiation of CSCs into SDCs significantly increased resistance to TMZ for all three lines (Fig 5B), despite downregulation of MGMT protein observed for HF2927 and HF3016 SDCs (Fig 4A).

**Fig 4.**
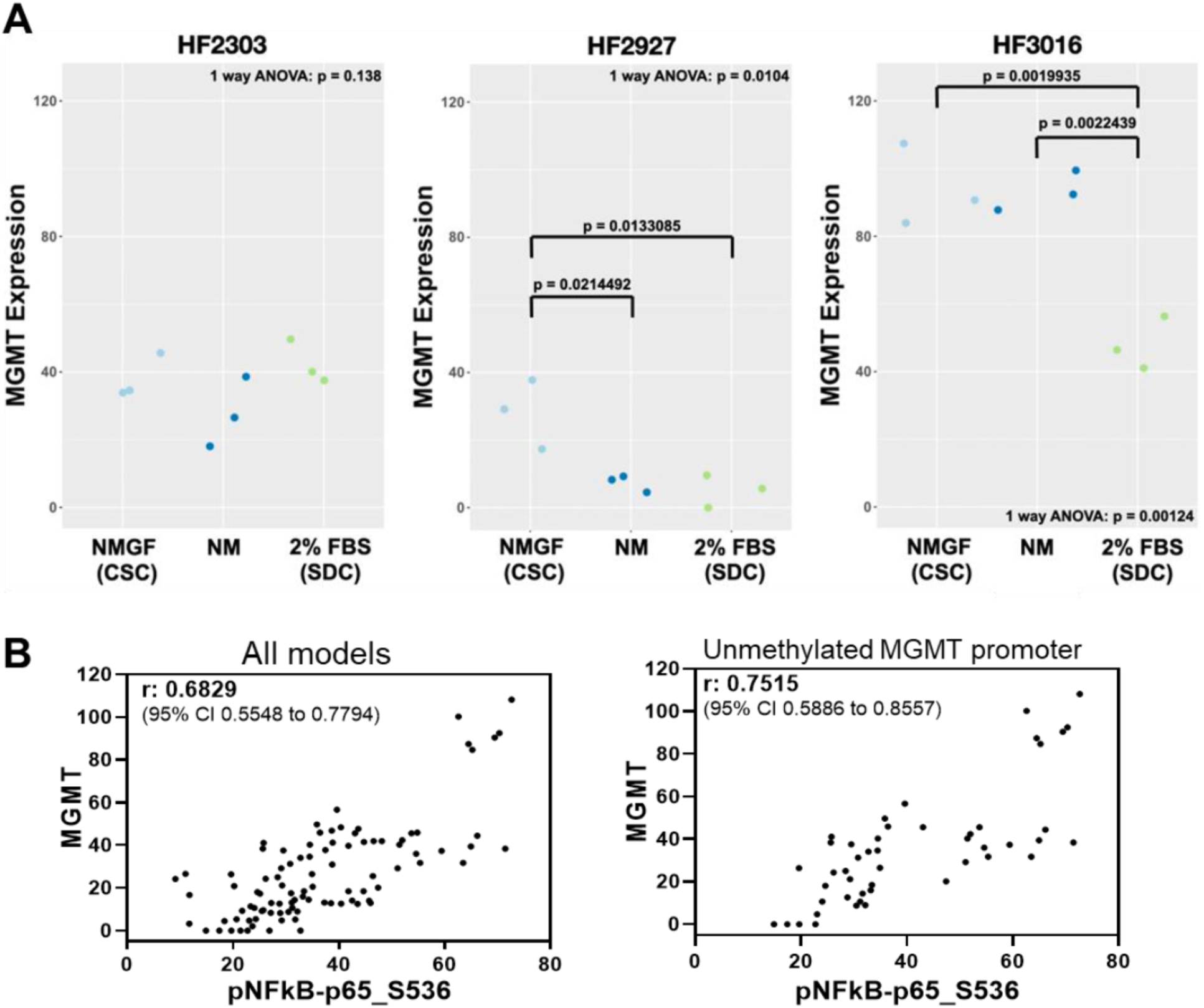
The impact of culture conditions on MGMT protein expression is cell line specific and correlates with NF-kB activation. **A)** MGMT protein levels measured by RPPA for HF2303, HF2927 and HF3016, in biological triplicates (S1 Table). P-values for 1-way ANOVA and post-hoc TukeyHSD test for comparison of the effect of culture conditions on MGMT protein expression are shown. Pairwise comparison between culture conditions was only significant for HF2927 and HF3016. **B)** Correlation of the levels of MGMT and phospho-NFkB (S536), for all 8 GBM models and culture conditions (left panel) and for the 4 models presenting unmethylated MGMT promoter (right panel). Spearman r and 95% CI are shown.

**Fig 5.**
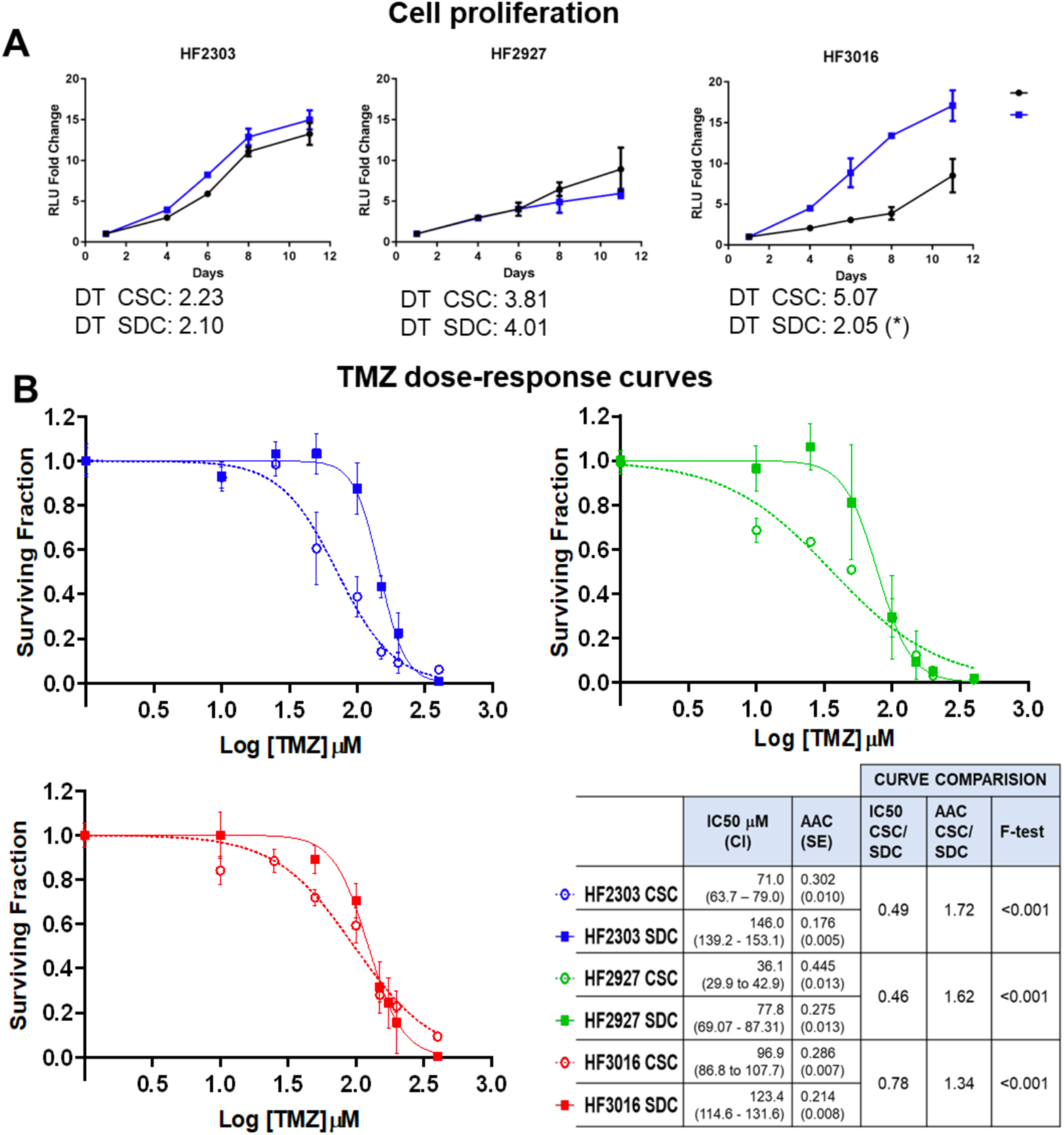
GBM CSCs are more sensitive to temozolomide than SDCs. **A)** To compare cell proliferation rates, CSCs and SDCs for each line were plated in the respective media in 96-well assay plates (1,000 cells/well) and cell viability was measured over 11 days in culture (n=5). Graphs represent mean (SE). The calculated doubling-times (DT, days) are shown, pairwise comparison between CSC and SDC values were significant for HF3016 only (*) p<0.01. **B)** Cells were plated in 96-well assay plates (2,000/plate) and treated for 5 days with the indicated doses of temozolomide, or equivalent DMSO control. Dose-response curves, the calculated IC50 concentrations with confidence interval (CI), and area above the curve (AAC) expressed as a fraction of total area with SE are shown. For each of the GBM patients, CSCs were significantly more sensitive to TMZ relative to SDCs, as shown by IC50 and AAC ratios, and p-values for dose-response curve comparison (F-test).

We then compared the sensitivity of CSCs and SDCs to clinically relevant RT doses. Isogenic populations of CSCs and their SDC progenies were exposed to 2 gray (Gy) and 4 Gy RT doses and cultured for 5 days, surviving fractions (SF2 and SF4) relative to control treatment were determined (Fig 6A) as previously described [51]. Sensitivity to RT was variable among these models, as a high degree of resistance was observed for HF2303 cells relative to both HF3016 and HF2927; HF2927 was the most sensitive (Fig 6A). On the other hand, no significant difference in sensitivity to RT between CSCs and SDCs were observed for these three models (Fig 6A). To evaluate the long-term effect of radiation, we compared the tumorigenic potential of control and RT treated CSCs. Cells expressing firefly luciferase (fLuc) were implanted intracranially in nude mice immediately after radiation (4 Gy), or mock radiation (control). Tumor growth was monitored by weekly bioluminescence imaging and by daily monitoring of symptoms associated with tumor burden. The three irradiated CSC lines exhibited a delay in tumor growth relative to controls, assessed by symptom-free survival, with all mice eventually developing tumors, demonstrating that cells surviving 4 Gy RT dose were tumorigenic. HF2303 displayed the highest resistance to RT in vitro and exhibited the least significant difference in the survival curves. On the other hand, the HF2927 cells, which were the most sensitive to RT in vitro, displayed the lowest p-value in the curve comparison (Fig 6B,C). The degree of difference in survival curves (Log-rank p-values) ranked according to the in vitro sensitivity of each cell line to RT.

**Fig 6.**
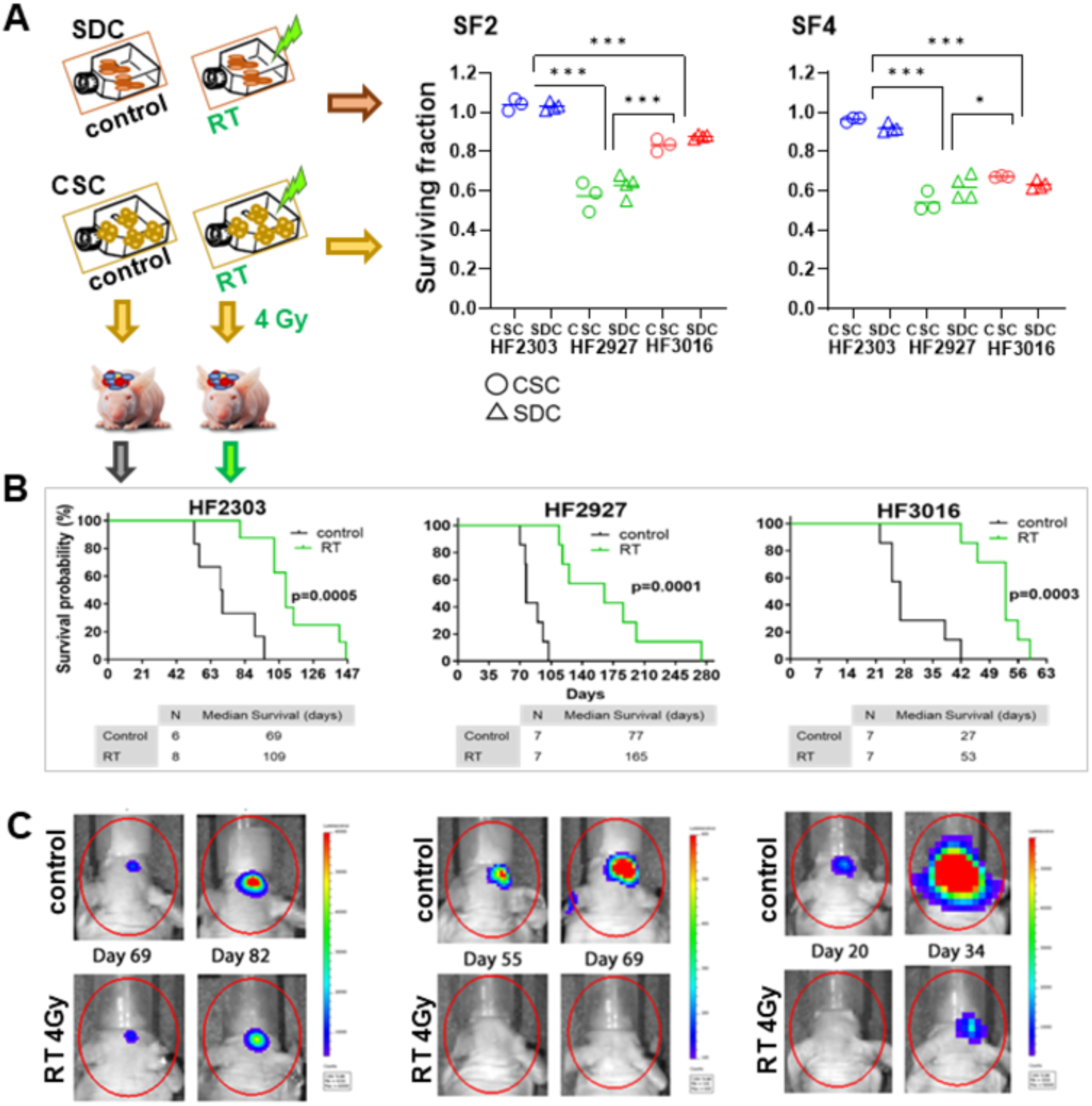
Response of glioblastoma CSCs and SDCs to ionizing radiation. **A)** CSC and SDC received a single radiation dose (2Gy or 4Gy) or mock radiation (control), and cell viability was measured 5 days later (n=3-4). No significant difference in sensitivity to RT between CSC and SDCs was observed for any of the lines (Holm-Sidak multiple comparisons test with adj p-value<0.05 threshold). The surviving fraction (SF) values among the 3 cell lines (CSC and SDC combined) were compared by one-way ANOVA, followed by Tukey’s multiple comparisons tests, adj p value<0.0001(***), <0.05 (*). **B)** Irradiated and control CSCs were implanted intracranially immediately after treatment, 3x10^5^ cells/mouse (n=6-8/group). Symptom-free survival for the orthotopic PDX is shown in Kaplan-Meier curves, compared by log-rank test. **C)** Longitudinal live bioluminescense images for representative PDX in (B).

### Transcriptional reprogramming in response to CSC differentiation

To determine how differentiation state impacts transcriptional reprogramming in response to TMZ and RT, we selected the most resistant (HF2303) and the most sensitive (HF2927) models. We first determined the pattern of global transcriptome changes after differentiation of CSCs into SDCs from bulk RNAseq data. To identify differentially expressed genes between the two differentiation states, we employed the non-parametric NOISeq R package, and subsequently performed pathway enrichment analysis, as detailed in the Materials and Methods section. Numerous signaling pathways and cellular processes enriched in CSCs were cell line specific. It has been proposed that CSCs, from GBMs and other cancers, tend to rely more on oxidative phosphorylation (OXPHOS) than the more differentiated cancer cells [52]. Here, we observed that OXPHOS gene expression signature was enriched in HF2303 CSCs but not HF2927 CSCs (Fig 7A). Interestingly, reserve respiratory capacity was higher in untreated HF2303 relative to HF2927 cells, and higher in SDC compared to CSCs (S3 Fig). Replication-dependent histones were highly enriched in HF2303 CSCs, in the absence of enrichment in cell proliferation markers (Fig 7A, S2 Table), as previously observed for embryonic and induced pluripotent stem cells [53]. Several nuclear pseudogenes with sequence similarity to mitochondrially encoded 16S rRNA (e.g. *MTRNR2L1, MTRNR2L3*), enriched in HF2303 CSCs (A in S2 Table) are predicted to encode humanin-like peptides and to be involved in negative regulation of the execution phase of apoptosis [54] (Fig 7A). IL2-STAT5 signaling was specifically upregulated in HF2927 CSC in relation to SDC (Fig 7A), in agreement with levels of pSTAT5_Y694 being elevated in HF2927 CSC (4.697 ± 0.98), while undetectable in HF2927 SDCs and HF2303 cells (B in S1 Table). Patterns of transcriptome reprogramming in response to differentiation that were common to both HF2927 and HF2303 models included downregulation of genes involved in cholesterol biosynthesis and upregulation of genes associated with ECM components and organization, including collagen type IV alpha 6 chain (*COL4A6*), a target of SRY-box transcription factor 2 (*SOX2*) transcriptional regulation [55] (Fig 7B). The expression of several transcription factors associated with stemness and pluripotency, such as *SOX2,* spalt like transcription factor 2 (*SALL2*) and POU class 5 homeobox 1 (*POU5F1* (Oct3/4)), was not altered upon differentiation of HF2303 and HF2927 CSCs into SDCs (A,B in S2 Table).On the other hand, other neural progenitor and stemness markers expressed in CSCs were down regulated upon differentiation: oligodendrocyte transcription factor 1 (*OLIG1*) in both lines; delta like canonical Notch ligand 3 (*DLL3*), integrin subunit alpha 6 (I*TGA6*), ETS variant transcription factor 4 (*ETV4*) in HF2927; fatty acid binding protein 7 (*FABP7*), oligodendrocyte transcription factor 2 (*OLIG2*) and ELOVL fatty acid elongase 2 (*ELOVL2*) [56] in HF2303 (Fig 7A, B).

**Fig 7.**
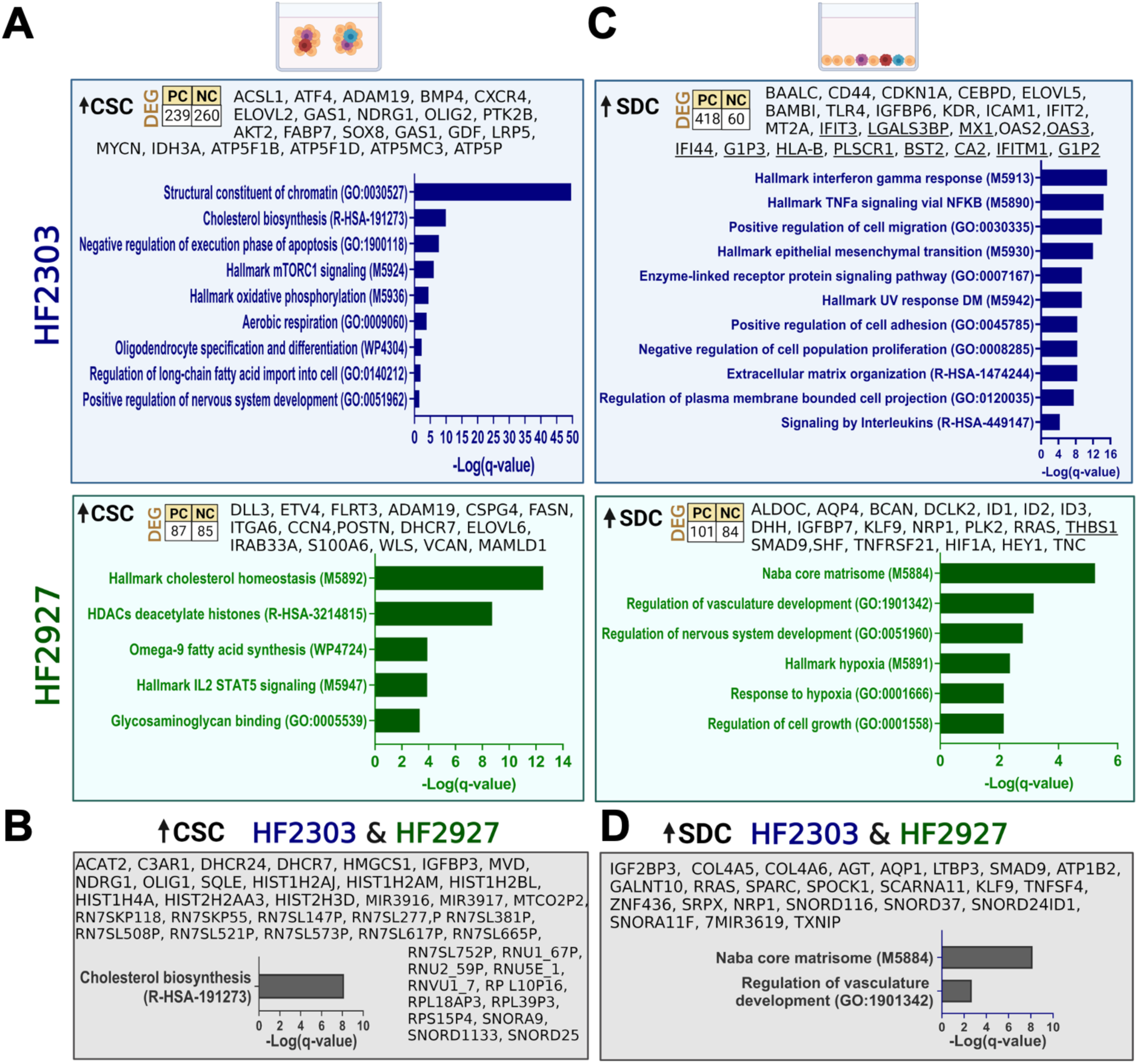
Transcriptional reprograming in glioblastoma cells in response to CSC differentiation. HF2927 and HF2303 CSC were grown in regular neurosphere media or in SDC cultures for 2 weeks. Total RNA from triplicate samples for each group was sequenced analyzed and DEG between CSC and SDC determined for each line. Total number of protein coding (PC) and non-coding (NC) genes, select genes and Metascape gene-set enrichment tests adjusted for multiple comparisons (q-value < 0.05) are shown for genes upregulated in CSCs (A) and in SDCs (C). The common DEGs for both lines along with enriched cellular processes are presented: 43 genes upregulated in CSCs (B) and 25 genes upregulated in SDCs (D).

IFN and interleukins signaling were specifically upregulated in HF2303 SDCs (Fig 7C). Indeed 29 genes associated with IFN response pathways were upregulated in HF2303 SDCs (Table 1), including members of the 49-gene interferon-related DNA damage resistance signature (IRDS) [57]. Differentiation of CSCs into SDCs resulted in enrichment of the epithelial mesenchymal transition (EMT) signature for HF2303 but not for HF2927 (Fig 7C), consistent with the transcriptional subclasses affiliation, mesenchymal for HF2303 and classical for HF2927 (Fig 1A). In agreement with our observation that receptor dependent SMAD1/3/8 are activated in GBM SDCs (Fig. 2A), inhibitor of DNA binding 1 (*ID1*), *ID2* and *ID3*, BMP/retinoic acid inducible neural specific 1 (*BRINP1*), downstream of BMP signaling, were upregulated in HF2927 SDCs (Fig 7C). Several components of the TGFb superfamily signaling were upregulated in HF2303 SDCs, including transforming growth factor beta receptor 2 (*TGFBR2*), *BMP1*, latent transforming growth factor beta binding protein 1 (*LTBP1*), *LTBP3*, *SMAD3*, *SMAD9* (Fig 7C,D; S2 Table). On the other hand, endogenous *BMP4* mRNA was downregulated and BMP antagonists DAN family BMP antagonist (*NBL1*) and BMP and activin membrane bound inhibitor (*BAMBI*) were upregulated in HF2303 SDCs, suggesting a possible negative feedback loop, specific for this model. BAALC binder of MAP3K1 and KLF4 (*BAALC)*, a regulator of developing neuroectoderm tissues overexpressed in acute leukemia and GBM [58], and a target of SOX2 in differentiated GBM cells [26], was upregulated in HF2303 SDCs (Fig 7C). Toll like receptor 4 (*TLR4*), which is expressed in both immune and neoplastic cells, was also upregulated in HF2303 SDCs (Fig 7C), in agreement with previous studies [59]. *HIF1A, an* IFN responsive gene (Table 1), along with downstream hypoxia markers were uniquely upregulated in HF2927 SDCs, under normal oxygen levels (Fig 7C)’. The 24 genes commonly upregulated in SDCs for both lines were enriched in ECM components (Fig 7D). Small nucleolar RNAs were also upregulated upon differentiation of both HF2303 and HF2927 (Fig 7D; S2 Table), including SNORD116 cluster, shown to have non-canonical functions in the differentiation, survival, and proliferation of neuronal cells [60].

**Table 1.** Differentially expressed genes associated with IFN response and signatures.

| Gene Symbol | DEG in response to 2% FBS differentiation |  | DEG in response to TMZ |  | DEG in response to RT |  | IFN response |
| --- | --- | --- | --- | --- | --- | --- | --- |
|  | Upregulated | Downregulated | Upregulated | Downregulated | Upregulated | Downregulated |  |
| *ACSL5* | - | - | HF2927 SDC | - | - | - | a |
| *ODF3B* | - | - | HF2927 SDC | - | - | - | a |
| *NEAT1* | - | - | HF2927 SDC | - | - | - | d |
| *SECTM1* | - | - | HF2927 SDC | - | - | - | c |
| *OASL* | - | - | HF2927 SDC;<br>HF2927 CSC | - | - | - | a, b, c, d |
| *CDKN1A* | HF2303 | - | HF2927 SDC;<br>HF2927 CSC | - | - | - | c |
| *RSAD2* | - | - | HF2927 SDC;<br>HF2303 SDC | - | - | - | a, b, c |
| *CMPK2* | - | - | HF2927 CSC | - | - | - | a, b, c |
| *CXCL10* | - | - | HF2927 CSC | - | - | - | a, b, c, d |
| *EPSTI1* | - | - | HF2927 CSC | - | - | - | a, b, c |
| *DHX58* | - | - | HF2927 CSC | - | - | - | a, b, c |
| ***MX2*** | - | - | HF2927 CSC | - | - | - | a, c, d |
| ***ISG15*** | HF2303 | - | HF2927 CSC | - | - | - | a, b, c, d |
| ***TRIM22*** | - | - | HF2927 SDC;<br>HF2927 CSC | - | HF2927 CSC | - | a |
| ***MX1*** | HF2303 | - | HF2927 CSC;<br>HF2303 CSC;<br>HF2303 SDC | - | HF2927 CSC | - | a, b, c, d, e |
| ***IFI6*** | HF2303 | - | HF2927 CSC | - | HF2927 CSC | - | a, d |
| ***OAS2*** | - | - | - | - | HF2927 CSC | - | a, c |
| ***HERC6*** | - | - | - | - | HF2927 CSC | - | a, b, c, d |
| ***IFI44L*** | - | - | - | - | HF2927 CSC | - | a, b, c, d |
| ***HLA-E*** | HF2303 | - | - | - | HF2927 CSC | - | a |
| ***OAS1*** | - | - | HF2303 SDC | - | - | - | a, b, d, e |
| ***OAS3*** | - | - | HF2303 SDC | - | - | - | a, c, d |
| ***SAMD9L*** | - | - | HF2303 SDC | - | - | - | a, b, c |
| ***GIMAP2*** | - | - | - | HF2927 SDC | - | - | a |
| ***TNFAIP3*** | - | - | - | HF2927 SDC | - | - | c |
| ***THBS1*** | - | HF2927 | - | HF2927 SDC | - | - | d |
| ***TNFAIP6*** | - | HF2927 | - | HF2927 SDC | - | - | c |
| ***HLA-B*** | HF2303 | - | - | - | - | HF2303 CSC | a, c, d |
| ***IGF2*** | - | - | - | - | - | HF2303 CSC | d |
| ***MTHFD2*** | - | HF2303 | - | - | - | - | c |
| ***IFIT3*** | HF2303 | - | - | - | - | - | a, b, c, d, e |
| ***BST2*** | HF2303 | - | - | - | - | - | a, b, c, d |
| ***IFI44*** | HF2303 | - | - | - | - | - | a, b, c, d, e |
| ***RIGI*** | HF2303 | - | - | - | - | - | a, c |
| ***IRF9*** | HF2303 | - | - | - | - | - | a, b, c |
| ***IFIT2*** | HF2303 | - | - | - | - | - | a, b, c |
| ***IFITM1*** | HF2303 | - | - | - | - | - | a, b, d |
| ***PLSCR1*** | HF2303 | - | - | - | - | - | a, b, c, d |
| ***PLAAT4*** | HF2303 | - | - | - | - | - | a |
| ***HLA-C*** | HF2303 | - | - | - | - | - | a, b |
| ***LGALS3BP*** | HF2303 | - | - | - | - | - | a, b, c, d |
| ***CA2*** | HF2303 | - | - | - | - | - | d |
| ***ARL4A*** | HF2303 | - | - | - | - | - | c |
| ***BTG1*** | HF2303 | - | - | - | - | - | c |
| ***C1R*** | HF2303 | - | - | - | - | - | c |
| ***CCL2*** | HF2303 | - | - | - | - | - | c |
| *FGL2* | HF2303 | - | - | - | - | - | c |
| *ICAM1* | HF2303 | - | - | - | - | - | c |
| *MVP* | HF2303 | - | - | - | - | - | c |
| *NAMPT* | HF2303 | - | - | - | - | - | c |
| *PELI1* | HF2303 | - | - | - | - | - | c |
| *UPP1* | HF2303 | - | - | - | - | - | c |
| *MT2A* | HF2303 | HF2927 | - | - | - | - | c |
| *HIF1A* | HF2927 | - | - | - | - | - | c |

(a) S3\_Table TCGA IFN response correlation group; (b) M5911 Hallmark IFN alpha response https://www.gsea-msigdb.org/; (c) M5913 Hallmark IFN gamma response https://www.gsea-msigdb.org/; (d) IRDS ref. [57]; (e) IFN/STAT1 survival proneural ref. [63]

### Transcriptional response to sublethal treatment with temozolomide and radiation is determined by genomics and differentiation state

To compare the transcriptional response to temozolomide treatment at distinct differentiation states, GBM cells were treated in triplicate with the respective TMZ IC40 concentrations, calculated from dose-response curves in Fig 5B: 58 mM (HF2303 CSC), 25 mM (HF2927 CSC), 134 mM (HF2303 SDC), 70 mM (HF2927 SDC), or equivalent DMSO control for 4 days. Cells were then harvested and total RNA isolated and sequenced. Differentially expressed genes between treated and control samples for each model and culture type were determined as described for Fig. 7 (C and D in S2 Table). Transcriptional response to 4-day TMZ treatment was largely regulated by wt p53 activation in HF2927 cells (Fig. 8A). We observed that 18 out of the 36 genes commonly upregulated in HF2927 CSCs and SDCs in response to TMZ treatment (Table 2) are direct transcriptional targets of p53 [55]. However, the differentiation state of HF2927 cells also influenced p53 transcriptional specificity, as 23 and 26 p53 target genes were significantly upregulated in response to TMZ exclusively in CSCs or SDCs, respectively (Table 2). In contrast, interleukin-27 (IL-27) mediated signaling pathway was enriched in HF2303 SDC in response to TMZ (Fig 8A), based on the upregulation of Myxovirus resistance dynamin like GTPase 1 (*MX1*), 2’ 5’ oligoadenylate synthetase 1 (*OAS1*) and *OAS2*, all of which are members of the IRDS [57], providing further evidence for the proposed cooperation between IL-27 and IFN signaling in response to DNA damage [61]. To identify other components of IFN signaling in GBMs, we identified 100 genes whose expression presented the highest correlation to *OAS1* (Spearman’s r > 0.55; q value < 5.0E-12) in The Cancer Genome Atlas (TCGA) GBM data set (CBIOPortal [62], accessed on 01/19/24). 62% of these genes are members of the hallmark IFN alpha or gamma response pathways, 24% are members of the IRDS [57], and this gene list also includes the IFN/STAT1 prognostic signature proposed for GBM proneural subtype [63] (S3 Table). We observed activation of IFN response in TMZ-treated HF2927 CSCs and SDCs, and in HF2303 SDCs (Fig 8A, Table 1). TMZ treatment induced expression of ECM components, and several neural developmental growth and survival factors, including brain derived neurotrophic factor (*BDNF*) and glial cell derived neurotrophic factor (*GDNF*), in HF2927 CSCs and SDCs (Fig 8A). Expression of other genes associated with neural development were down regulated in TMZ treated HF2927: platelet derived growth factor receptor alpha (*PDGFRA*), *OLIG2*, and prominin 1 (*PROM1*) in SDCs; neuropilin 1 (*NRP1*), GLI family zinc finger 1 (*GLI1*), and Kruppel like factor 9 (*KLF9*) in CSCs (Fig 8A). *CDKN1A* (p21), the main effector of p53-mediated downregulation of cell cycle genes, leading to cell cycle arrest, was upregulated in response to TMZ in both HF2927 CSCs and SDCs (Fig 8A,D). The more pronounced downregulation of cell cycle genes in HF2927 SDCs relative to CSCs (Fig 8A) is consistent with corresponding upregulation of Lin-37 DREAM MuvB core complex component (*LIN37*)in SDC but not in CSC (Fig 8A), as LIN37 is required for transcriptional repression by the DREAM complex, downstream of p53 signaling [64].A possible explanation for these observations is that cell cycle-arrested TMZ-treated HF2927 CSCs may have undergone apoptosis to a greater extent at the time point analyzed, as they demonstrated increased sensitivity to TMZ treatment (Fig 5B). Furthermore, genes coding for proteins involved in EMT, including the master regulator twist family bHLH transcription factor 1 (*TWIST1*), were upregulated in response to TMZ exclusively in HF2927 SDCs, which could contribute to the increased resistance of the differentiated cells.

**Fig 8.**
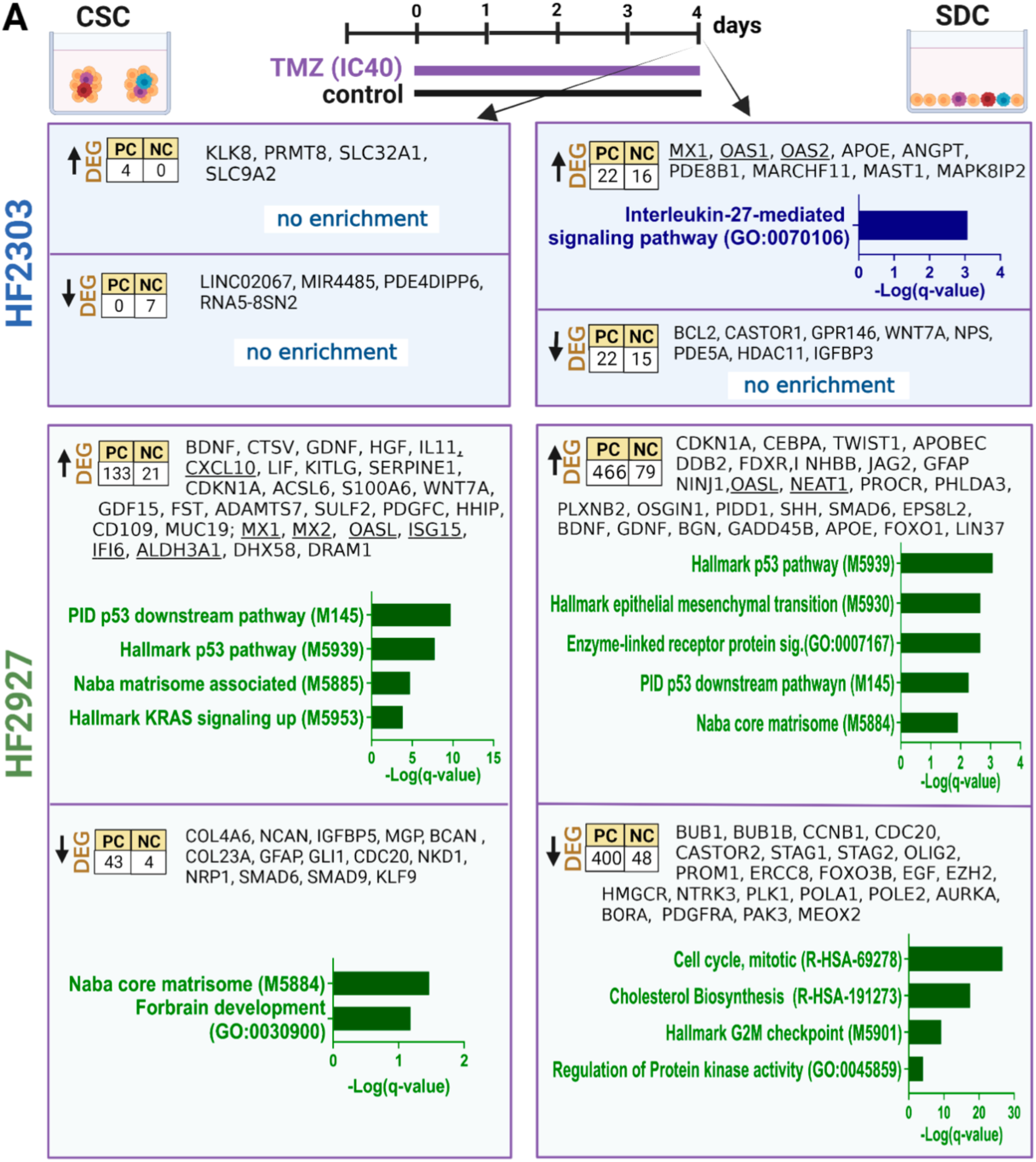
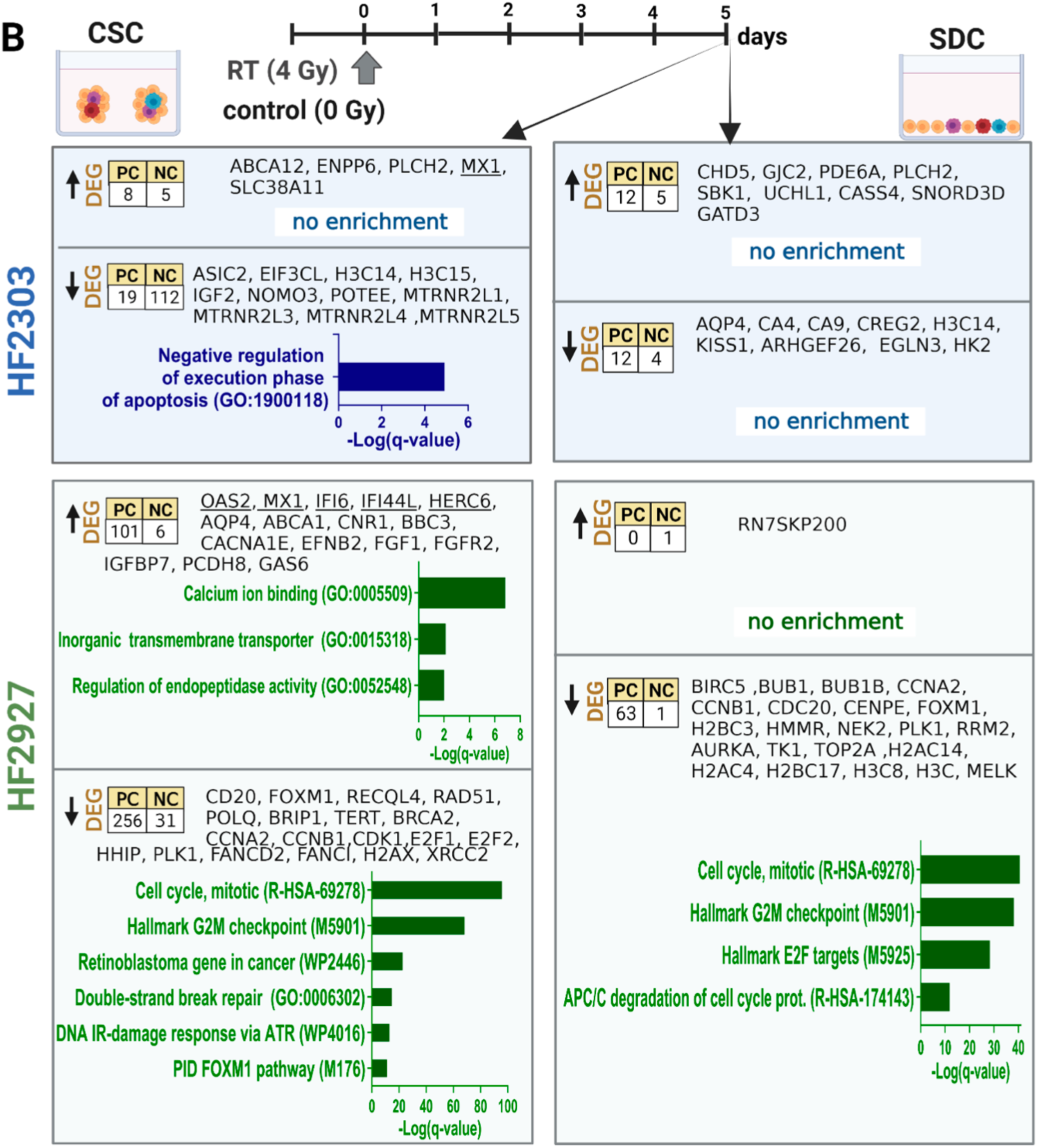
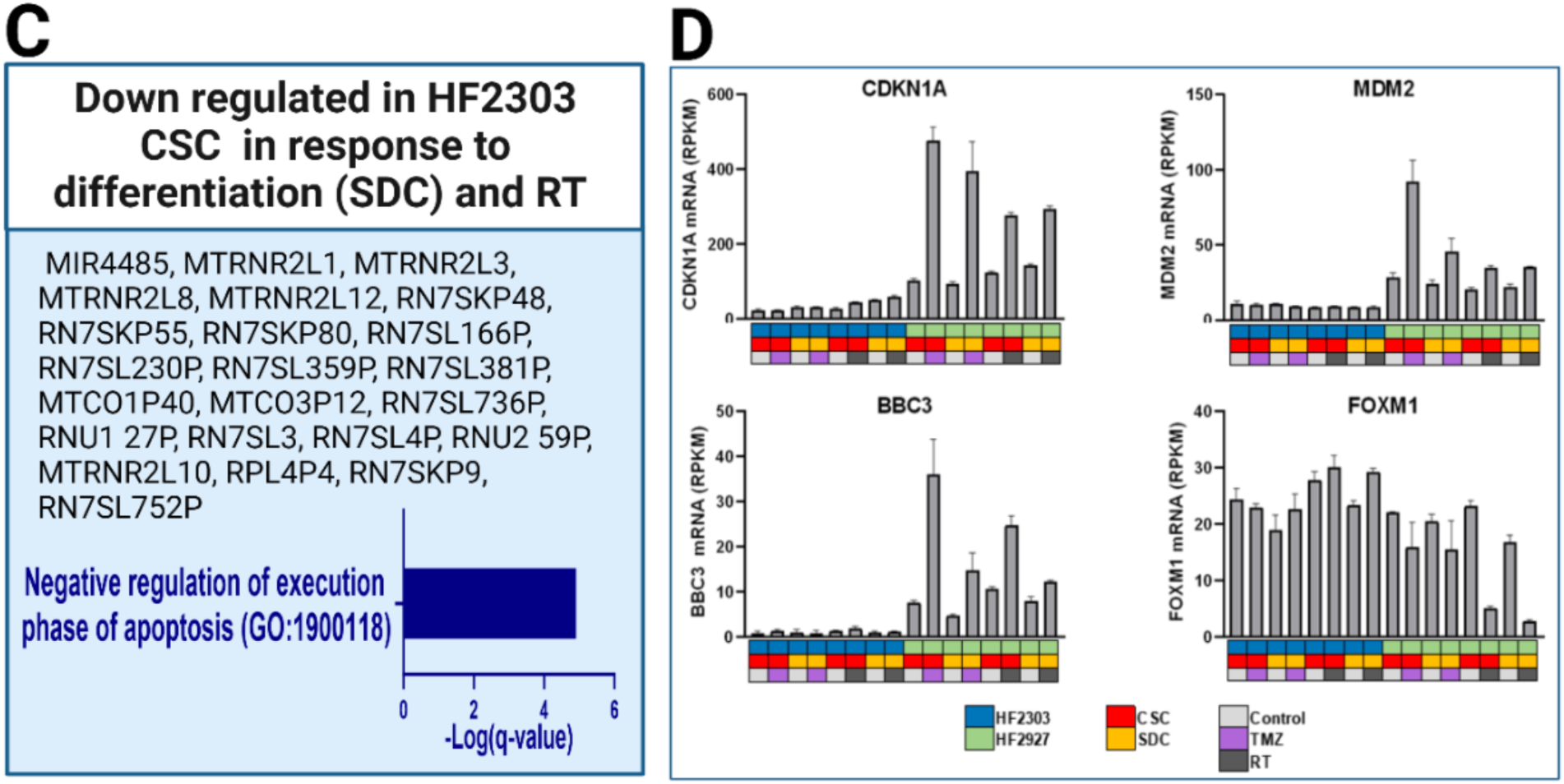
Transcriptional response to genotoxic treatment in glioblastoma cells. Differentially expressed genes between treated and control samples were determined and enrichment performed as described for Fig. 7. **A)** Transcriptional changes in CSCs and SDCs in response to 4-day TMZ treatment (IC40 concentrations). **B)** Transcriptional changes in CSCs and SDCs measured 5 days after one dose RT (4 Gy). **C)** RNA commonly downregulated in HF2303 CSCs in response to differentiation and RT treatment. **D)** mRNA expression (RPKM) of p53 transcriptional targets CDKN1A (p21), MDM2, and BBC3, and of cell cycle regulators in HF2303 and HF2927 control and treated cells, mean and SE of n=3. Figures created with Biorender.com.

**Table 2.**
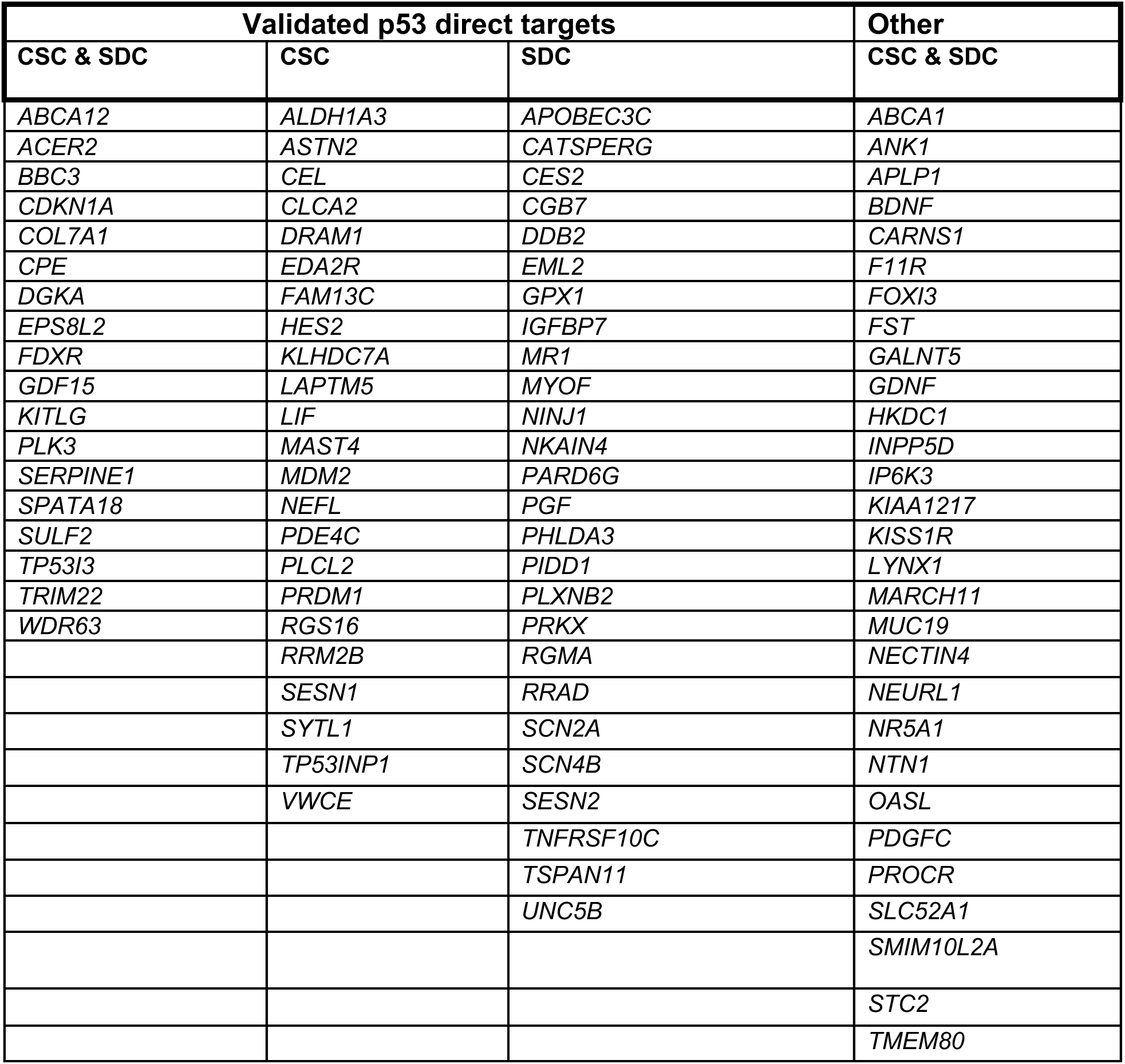
Genes upregulated in HF2927 cells in response to temozolomide.

| **Validated p53 direct targets** |  |  | **Other** |
| --- | --- | --- | --- |
| **CSC & SDC** | **CSC** | **SDC** | **CSC & SDC** |
| *ABCA12* | *ALDH1A3* | *APOBEC3C* | *ABCA1* |
| *ACER2* | *ASTN2* | *CATSPERG* | *ANK1* |
| *BBC3* | *CEL* | *CES2* | *APLP1* |
| *CDKN1A* | *CLCA2* | *CGB7* | *BDNF* |
| *COL7A1* | *DRAM1* | *DDB2* | *CARNS1* |
| *CPE* | *EDA2R* | *EML2* | *F11R* |
| *DGKA* | *FAM13C* | *GPX1* | *FOXI3* |
| *EPS8L2* | *HES2* | *IGFBP7* | *FST* |
| *FDXR* | *KLHDC7A* | *MR1* | *GALNT5* |
| *GDF15* | *LAPTM5* | *MYOF* | *GDNF* |
| *KITLG* | *LIF* | *NINJ1* | *HKDC1* |
| *PLK3* | *MAST4* | *NKAIN4* | *INPP5D* |
| *SERPINE1* | *MDM2* | *PARD6G* | *IP6K3* |
| *SPATA18* | *NEFL* | *PGF* | *KIAA1217* |
| *SULF2* | *PDE4C* | *PHLDA3* | *KISS1R* |
| *TP53I3* | *PLCL2* | *PIDD1* | *LYNX1* |
| *TRIM22* | *PRDM1* | *PLXNB2* | *MARCH11* |
| *WDR63* | *RGS16* | *PRKX* | *MUC19* |
|  | *RRM2B* | *RGMA* | *NECTIN4* |
|  | *SESN1* | *RRAD* | *NEURL1* |
|  | *SYTL1* | *SCN2A* | *NR5A1* |
|  | *TP53INP1* | *SCN4B* | *NTN1* |
|  | *VWCE* | *SESN2* | *OASL* |
|  |  | *TNFRSF10C* | *PDGFC* |
|  |  | *TSPAN11* | *PROCR* |
|  |  | *UNC5B* | *SLC52A1* |
|  |  |  | *SMIM10L2A* |
|  |  |  | *STC2* |
|  |  |  | *TMEM80* |

Ionizing radiation can damage molecules directly, exerting most of its toxicity to cancer cells through DNA double strand breaks (DSBs), and indirectly through the generation of free radicals. The complex cellular responses leading to cell cycle arrest, apoptosis, DNA repair and other processes occur over time [65]. The relative sensitivity of glioblastoma cells to RT varied according to tumor of origin but not according to differentiation status under the experimental parameters employed here (Fig 6A). Furthermore, pre-implantation treatment of CSCs with 4Gy led to delayed xenograft tumor growth for all models (Fig 6B). To integrate these observations with cellular responses, CSCs and SDCs were treated in triplicates with 4 Gy radiation dose or mock radiation control, followed by incubation in the respective growth media for 5 days prior to RNA isolation and sequencing. Under these conditions, few transcripts were altered in response to RT in HF2303 (Fig 8B), consistent with the resistance to 4 Gy dose observed for both SDCs and CSCs (Fig 6A). The only transcripts commonly upregulated in HF2303 CSCs and SDCs in response to RT were phospholipase C eta 2 (*PLCH2*) and *SNORD3D* (Fig 8B). Interestingly, 50 of the pseudogenes involved in negative regulation of the execution phase of apoptosis, which were downregulated in response to differentiation of CSCs into SDCs (Fig 7), were also downregulated in RT-treated HF2303 CSCs (Fig 8B,C). Compared to TMZ treatment, fewer IFN response genes were upregulated in RT treated cells, including *MX1* which was upregulated in both CSC lines, and *OAS2*, *HERC6*, *IFI44L* and *HLA-E* in HF2927 CSCs (Table 1, Fig 8B). In contrast to the response to TMZ, where the cells were analyzed after 4 days of continuous treatment (Fig 8A), p53-mediated signaling in HF2927 was more attenuated 5 days after RT, as expected (Fig 8B). Few p53 transcriptional targets remained elevated in irradiated HF2927 cells, such as BCL2 Binding Component 3 (*BBC3 / PUMA*), a pro-apoptotic BCL-2 family member, while expression of several others, including *MDM2*, were not altered at the 5-day timepoint (Fig 8B,D). *CDKN1A*/p21, the main effector of p53 mediated cell cycle arrest significantly upregulated in response to TMZ (Fig 8A), was elevated to a lesser degree in response to RT, below the statistical threshold employed to determine DEGs (Fig 8D). However, sustained downregulation of cell cycle genes downstream of p53 activation was observed for RT-treated HF2927 CSCs and SDCs (Fig 8B). Indeed, 53 out of the 56 transcripts commonly downregulated in HF2927 CSC and SDCs in response to RT are transcriptional targets of the DREAM (dimerization partner, RB-like, E2F and multivulval class B) repressive complex [66] (Table 3). Among these targets, forkhead box M1 (*FOXM1*), a transcriptional activator of cell cycle genes, especially those involved in G2 and M phases [67], along with its transcriptional targets, including Polo-like kinase 1 (*PLK1*) which phosphorylates and activates FOXM1 in a positive feed-back loop [68], were downregulated exclusively in RT-treated cells at the analyzed time point (Fig. 8B,D). Furthermore, DNA damage response and double strand break repair genes were downregulated in response to RT predominantly in HF2927 CSCs (Fig 8B), indicating that these cells would be sensitized to subsequent DNA damaging interventions.

**Table 3.**
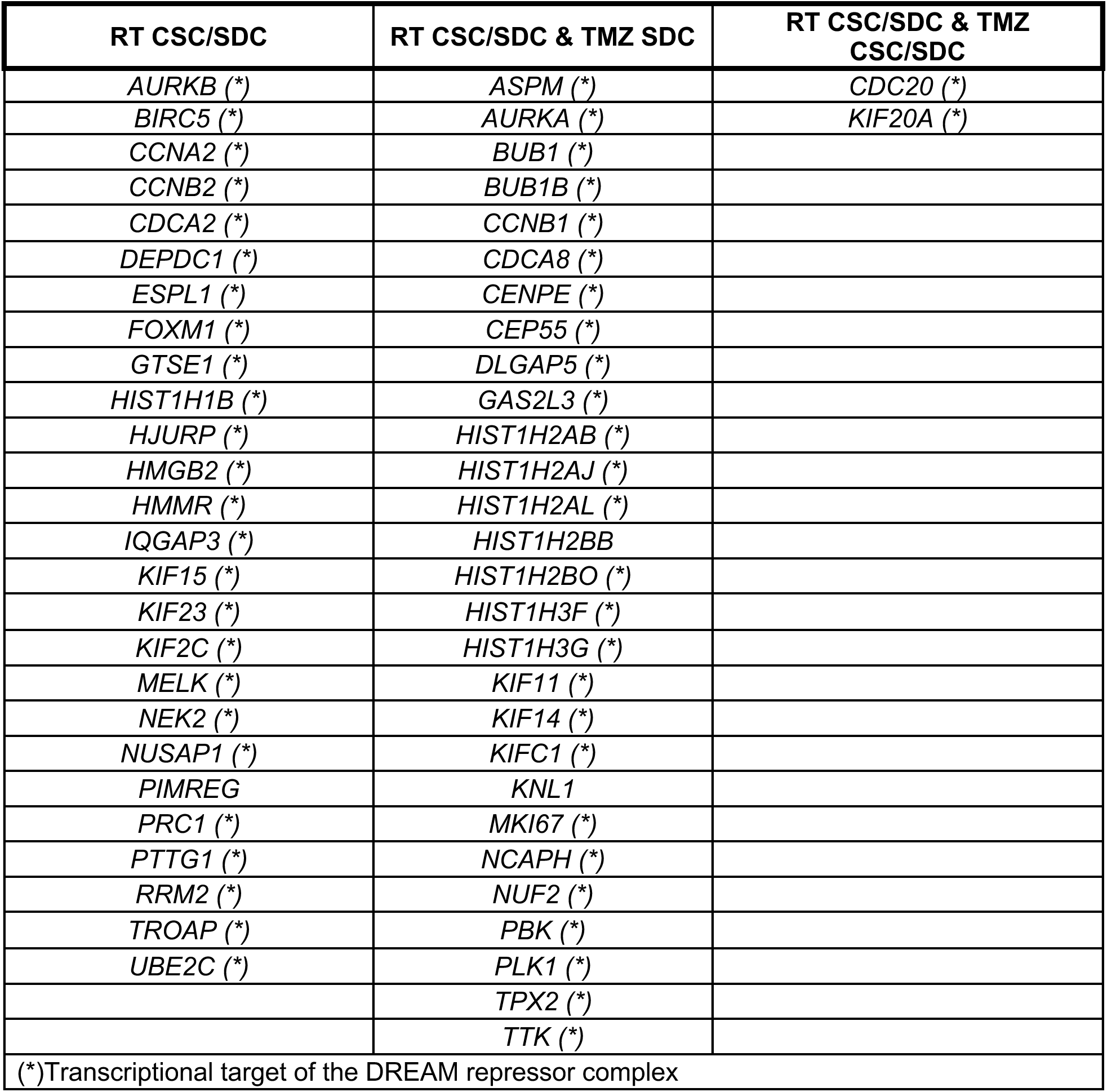
Genes downregulated in HF2927 CSC and SDC in response to radiation.

### DNA methylation alteration pattern in differentiation of CSCs

To verify the contribution of epigenetic regulation to the transcriptional responses reported above, we analyzed the global DNA methylation profile of control and treated HF2303 and HF2927 CSCs and SDCs in triplicates, using Infinium Human Methylation 450K BeadChip. Differences in DNA methylation distinguished the samples according to the tumor of origin (HF2303 vs HF2927), as shown in the t-distributed stochastic neighbor embedding (t-SNE) plot (Fig 9A). Analysis of the differentially methylated positions (DMPs) and regions (DMRs) between CSC and SDC for each of these models identified 892 DMP and 12 DMRs for HF2303, and 27,431 DMP and 2 DMRs for HF2927 (S4 Table). Similar to the significant overlap in DEGs associated with CSC differentiation between the two cell lines (Fig 7B, D, Fig 9B), we identified 384 DMPs common to both lines (Fig 9B, S4 Table), with a pattern of decreased b-values in SDCs relative to CSCs in HF2303, while HF2927 SDCs exhibited an increase in b-values. The intersection between genes mapped to the DMPs, at the transcription start sites or within CpG islands, and the genes differentially expressed between CSCs and SDCs were limited to 17 and 59 genes for HF2303 and HF2927, respectively. Among these genes, latent transforming growth factor beta binding protein 3 (*LTBP3*) was associated with DMPs and upregulation in SDC for both models (Fig 9B, S4 Table). Cholesterol homeostasis, hypoxia, growth factor binding, and ECM components were significantly represented in the 59 gene list from HF2927 (S4 Table). No significant DNA methylation changes were observed in the acute response to RT and TMZ treatment across both models and differentiation states (S4 Fig).

**Fig 9.**
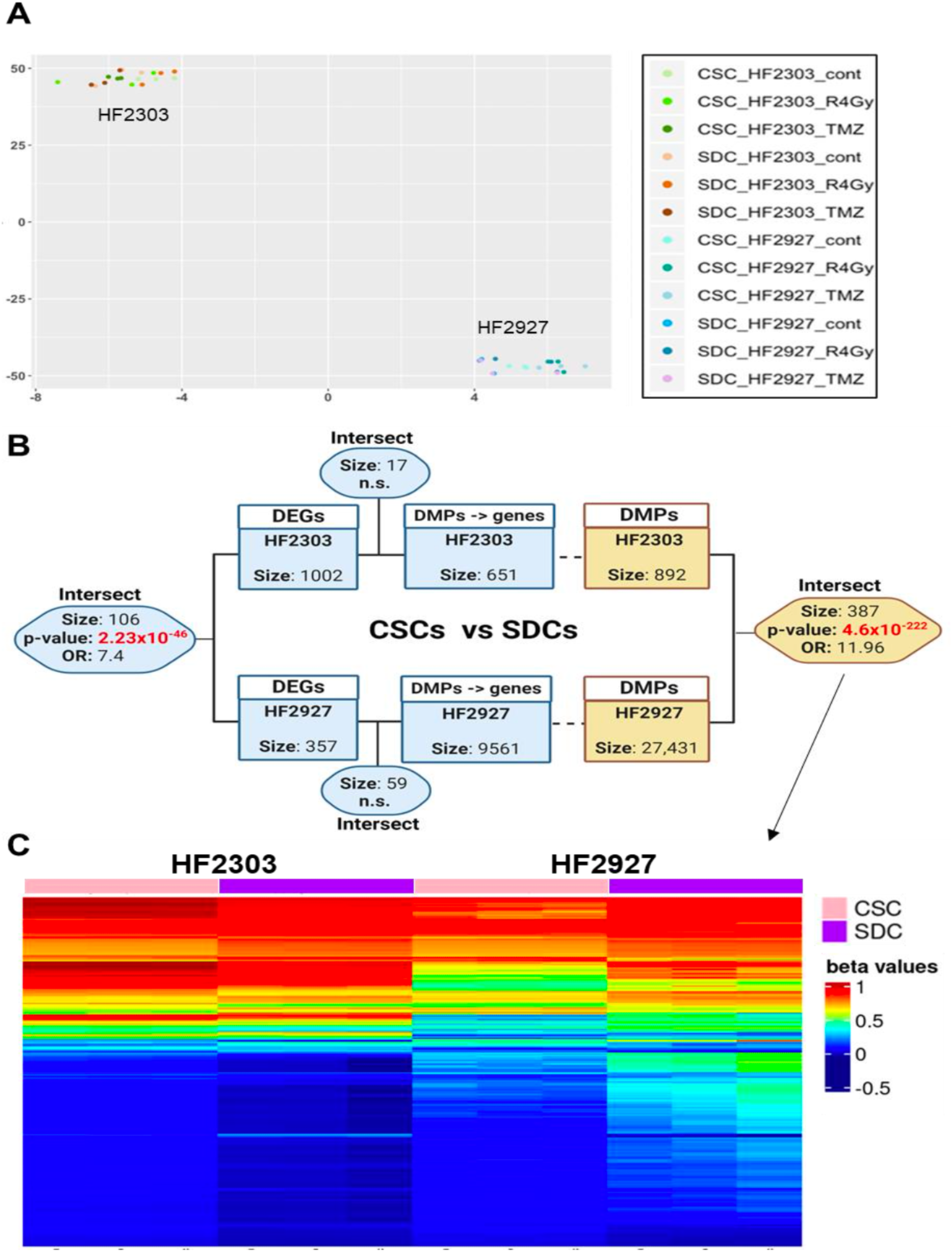
Genome-wide DNA methylation profiling groups samples by patient of origin and identifies DNA methylation alteration patterns between glioblastoma CSCs and their SDCs progeny. **A)** t-distributed stochastic neighbor embedding (t-SNE) plot from b-values for all CpG sites comparing HF2303 and HF2927 CSCs and SDCs treated with control, RT or TMZ, in triplicates. **B)** Schematic showing the significant intersect for HF2303 and HF2927 of differentially expressed genes (DEGs) and differentially methylated positions (DMPs) between CSCs and SDCs. The intersect between DMPs mapped to genes (see Methods) and DEGs are shown. Significance of the intersections was determined using GeneOverlap, p-values and estimated odds ratio are shown; n.s., nonsignificant. **C)** Heatmap showing the b-values for the 387 DMPs common to HF2303 and HF2927. Figure 9B was created with Biorender.com.

## Discussion

Understanding the extent to which oncogenic signaling and biological pathways modulating CSC phenotype and adaptation to external stimuli - such as astrocytic differentiation cues or growth factor withdrawal - are influenced by diverse genomic abnormalities, or are conversely generalizable to all GBMs, is crucial for the design and interpretation of biological and experimental therapeutical studies. For example, astrocytic differentiation of CSCs into SDCs resulted in the activation of RTK and MAPK pathways in most GBM cells, consistent with a role for MAPK in early astrocytic differentiation [69]. However, this activation was not observed for the two models carrying ecDNA EGFR amplification (HF2927 and HF3016). On the other hand, BMP-Smad pathway was prominently activated in all GBM models upon exposure to 2% and 10% FBS. Activation of BMP-Smad was further supported by the observation that transcriptional targets of r-Smad1/5/8 were upregulated in HF2303 and HF2927 SDCs. This is consistent with the use of BMP4 to induce the differentiation of iPSC into astrocytes [30], with BMP signaling inducing astrocytic differentiation of patient-derived oligodendroglioma cells [70], and with BMP ligands being present in FBS [33]. We found that signaling promoting cell survival in untreated GBM cells varies among the different models but not as much between CSCs and SDCs. For example, hyperactivation of the PI3K/AKT/mTOR pathway, which promotes cell growth, proliferation and survival, is a common feature of GBMs [71], was observed in both CSCs and SDCs for 7/8 GBM models, with a noted decrease in activation upon withdrawal of growth factors, as expected. The integrated analysis of targeted proteomics in multiple GBM models, distinct differentiation states and treatment groups showed a complex co-activation and cross talk among oncogenic signaling pathways, which greatly contributes to aberrant cancer cell growth, proliferation, survival, and resistance to therapeutic interventions [24]. We show that culturing CSCs in media supplemented with 10% FBS resulted in suppression of key oncogenic signaling in all 8 GBM models, providing further evidence that this media is not conducive to preserving GBM molecular characteristics, as previously reported [28, 47].

Earlier studies characterized GBM “CSCs” as a subpopulation of cells isolated from dissociated tumors based on expression of certain markers (e.g., CD133), which were subsequently cultured in neural stem cell media, while marker-negative cells were considered non-CSCs, or “differentiated”, and cultured in 10% FBS. Using this paradigm, these earlier studies attributed GBM marker positive “CSCs” with increased DNA repair capacity and corresponding resistance to RT, when compared to marker negative “non-CSCs”, e.g. [72]. In contrast, here GBM CSC subpopulations are propagated through enrichment in selective neurosphere media, a widely employed strategy [11, 20, 21], and differentiated GBM cells are defined as the progeny from astrocytic differentiation of CSCs in 2% FBS (SDCs). In this context, we report that sensitivity to clinically relevant doses of RT was unchanged between CSCs and their differentiated SDCs progeny, while SDCs presented increased resistance to TMZ for all three models tested, two representing unmethylated and one methylated MGMT promoter. We show that baseline levels of MGMT protein are strongly correlated with the levels of NF-kB activation in GBM cells, regardless of MGMT promoter methylation status, supporting previous reports that NF-kB activation confers resistance to DNA-alkylating agents in part through regulation of MGMT in glioma [7].

Transcriptional reprograming associated with the differentiation of CSCs into SDCs revealed two commonly altered cellular functions in the GBM models analyzed. Cholesterol biosynthesis was enriched in CSCs, with evidence for transcriptional activity of sterol regulatory element binding proteins (SREBPs), in response to the lack of cholesterol in the serum-free neurosphere media. Because GBM cells tend to uptake cholesterol from the brain microenvironment [73] and cholesterol is a component of FBS, cholesterol biosynthesis was downregulated in SDCs (2% FBS). ECM components, receptors and regulators were upregulated in SDCs, resembling transcriptional regulation of ECM components in astrocytes [74], likely with contribution of stimulatory attachment factors in FBS. The enrichment in ECM-receptor interaction observed for SDCs is associated with treatment resistance GBMs [75], and could be a factor contributing to the increased resistance of GBM SDCs to TMZ treatment. The retention of the expression of key stemness signature genes in SDCs, including *SOX2* and several of its transcriptional targets, highlights the role of these genes in the maintenance of developmental plasticity in more differentiated GBM cells, as we previously reported [26]. Other aspects of the transcriptional reprograming associated with differentiation of CSCs were model specific. We had previously shown that NF1 mutant HF2303 CSCs, like its parental tumor, belongs to the mesenchymal subclass and can recapitulate the biphasic phenotype associated with gliosarcoma, a GBM morphological variant, in orthotopic mouse xenografts [26]. Here, we show that CSC differentiation for this model resulted in enrichment of EMT signature, analogous to the mesenchymal cell state in NF1-deficient GBM [12]. DNA methylation alterations associated with CSC differentiation varied, with 30-fold more DMPs identified for HF2927 relative to HF2303, but the intersect between the 2 lines was highly significant at CpG level, but not at DMR level. On the other hand, under the experimental conditions employed in this study, we did not observe significant changes in global DNA methylation in response to treatment. There is no consensus in the occurrence and extent of changes in global DNA methylation patterns in response to RT or TMZ treatment in cultured cells [76]. This is in part due to the variability in cell intrinsic factors, such as cell type, genomic alterations, proliferation rate and cell cycle phase, as well as extrinsic factors, including culture conditions, RT and TMZ dosage and schedule.

We observed that both genomics and cell developmental states impact GBM response to TMZ and RT. The status of *TP53*, a key regulator of cellular response to DNA-damaging agents, was not surprisingly a key determinant of the transcriptional response of the GBM models to TMZ and RT. HF2927 carries wt TP53 while HF2303 carries TP53 G245S, a dominant negative mutation which disrupts the structure of p53 DNA binding domain, impairing p53 transcriptional activity [77]. Furthermore, wt p53 transcriptional activation was modulated by differentiation status, most notably in TMZ treated cells. Downregulation of cell cycle genes by p53 in response to DNA-damage, accomplished indirectly through the DREAM repressive complex [66], was observed for wt p53 CSCs and SDCs in response to TMZ and RT. *FOXM1,* a transcriptional activator of cell cycle genes overexpressed in various cancers [67], and a DREAM target [66], was highly expressed in untreated HF2303 and HF2927 cells. Here we show that *FOXM1*, along with its downstream signaling, were exclusively downregulated in HF2927 CSCs and SDCs 5 days after RT, although p53 target genes, including *CDKN1A*/p21 were no longer upregulated at this timepoint, indicating the prolonged effect of cell cycle arrest through downregulation of cell cycle genes, downstream of p53 activation, but we show these cells retained their tumorigenic potential.

Our work provides further evidence for a role of IFN response in resistance of GBMs to the standard of care. It is well stablished that accumulation of cytoplasmic ssDNA or dsDNA in cancer cells following DNA damage activates cGMP-AMP synthase (cGAS)–stimulator of IFN genes (STING) pathway, leading to the activation of type I IFN production and upregulation of IFN-stimulated genes (ISGs) [78]. Conflicting roles for IFN response ISGs in malignancy and resistance to treatment have been proposed. For example, ISGs can promote RT resistance through suppression of IFN production, in a negative feedback mechanism [79], or sensitize p53-deficient cells to chemotherapy [80]. Others have shown that STAT1 suppression in classical GBM CSCs leads to chemo-resistance [81]. IRDS is a subset of ISGs induced by prolonged exposure to low doses of type I IFN [82], and associated with RT resistance to radiation in various cancers [57]. *OAS1,* a member of the IRDS [57] involved in resistance to DNA-alkylating therapy [83], is also part of the IFN/STAT1 prognostic signature proposed for GBM proneural subtype [63]. Because IFN response is coordinated and typically presents a high degree of correlation in gene expression, our analysis identified genes highly correlated with expression to *OAS1* in the TCGA GBM dataset. Upregulation of these IFN responsive genes was overrepresented in response to TMZ and RT treatment of GBM CSCs and SDCs. Differentiation of HF2303 CSCs was characterized by upregulation of IFN response genes. This is consistent with reports that cancer cell-intrinsic type I IFN signaling is inversely correlated to stemness in a pan-cancer study [84]. We did not observe this IFN activation in HF2927 SDCs cells, highlighting the heterogeneity of responses of GBM CSCs to external stimuli. Our results highlight the importance of the IFN pathway activation in modulating glioma response to the standard of care, warranting further exploration as a therapeutic target and predictive biomarker.

## Conclusions

These results from the integrated analysis of treatment response and differentiation in bulk patient-derived GBM cell populations, which reflect the cumulative dynamic alterations in CSC and SDC sub-compartments, provide both validation of existing research and novel insights for further in vivo investigation, considering the added complexity of interactions among neoplastic cells in different states and the tumor microenvironment. Our findings have important implications for pre-clinical studies, highlighting the need for caution when generalizing results related to the differentiation of CSCs or their treatment sensitivity and molecular response across diverse GBM models.

## Methods

### Cell culture

Glioblastoma patient-derived cancer stem cell cultures were obtained from the live biobank collection at the Hermelin Brain Tumor Center, Henry Ford Hospital (Detroit, MI). These cells were derived from de-identified surgically resected glioblastoma specimens, collected with written informed consent, under a protocol approved by the HFH Institutional Review Board as previously described [85], in compliance with all relevant ethical regulations for research using human specimens. Cells were cultured in serum-free neurosphere media consisting of Dulbecco’s Modified Eagle Medium (DMEM)/F12 media (Invitrogen), N2 supplement (Gibco) and 0.5 mg/ml BSA, supplemented with growth factors 20 ng/ml EGF and 20 ng/ml bFGF (Peprotech) (NMGF), to select for CSCs [85]. CSC cultures were passaged in vitro and used prior to achieving passage 20. Identity of CSC lines was confirmed by comparing genotype with the patient germline using Short Tandem Repeat (STR) analysis. CSCs were dissociated and cultured for 2 weeks in three different conditions: NM lacking growth factors, and NMGF supplemented with 2% or 10% FBS (HyClone). Neural progenitor cells (NPCs) were derived from H9 human ESC, obtained from WiCell (Madison WI) as described [86] and differentiated in NMGF supplemented in 2% FBS [87].

### Reverse Phase Protein Arrays analysis

Cell lysates were obtained in triplicate for each cell line and growth conditions described (Fig 1B). The levels of 66 proteins/post-translational modifications in the cell lysates were analyzed by Reverse Phase Protein Arrays (RPPA) at the Mason’s Center for Applied Proteomics and Molecular Medicine (George Mason University), essentially as described previously [31, 32]. Statistical analyses of RPPA normalized values were conducted in the R statistical environment (4.2.1). Heatmaps were generated by ComplexHeatmap package in R (2.12.1) [88]. Z-scores for each protein and post-translational modification were calculated based on population mean and standard deviation ((x-μ)/σ). Fold-change was calculated using the gTools package (3.9.3) and computed as follows: num/denom if num>denom, and as -denom/num otherwise. Dendrograms were produced using Spearman correlation as the distance method.

### Cell proliferation, radiation and temozolomide treatment

#### Cell proliferation

Cell proliferation curves were established by plating 1,000 cells/well into 96-well plates in NMGF (CSCs) or NMGF supplemented with 2% FBS (SDCs) and measuring cell viability after 1, 4, 6, 8, and 11 days in culture (n=5). Wells were supplemented with fresh media every 3-4 days. Cell viability was measured using CellTiter-Glo Luminescent Cell Viability Assay (Promega), relative light units (RLU) were measured in a BioTek Cytation 3 plate reader. Doubling time (DT) was calculated using the formula DT=(t1-t2) ln2 / ln(RLU2/RLU1).

#### Temozolomide dose-response curves

CSCs and SDCs were plated in 96-well assay plates (3,000 cells/well), in quintuplicates and treated with 10 to 400 mM temozolomide (Schering-Plough) or DMSO control for 4 days. CellTiterGlo (Promega) was used to measure cell viability and dose-response curves were generated by non-linear fitting, IC50 doses and area above the curve (AAC) were calculated in GraphPad Prism (v.9.0).

#### Ionizing radiation treatment (RT)

CSC and SDC cells were dissociated, irradiated with 0, 2 or 4 Gy using a 5000 Ci Cesium (Cs-137) irradiator, and plated in triplicate at density of 3,000 cells/well in 96-well assay plates in the respective growth media and incubated under standard conditions for 5 days. Cell viability was assayed with CellTiter Glo (Promega). Surviving fractions at the two radiation doses were calculated and compared.

#### Tumorigenic potential of irradiated cells

Mouse experiments were performed under an approved Henry Ford Health Institutional Animal Care and Use Committee protocol (IACUC #1449) by properly trained personnel. Single cell suspensions of glioblastoma CSCs constitutively expressing firefly luciferase were irradiated with 0 (mock control) or 4Gy, as described above. Immediately following radiation or control treatment, 2x10^5^ cells/mouse were implanted intracranially in female NCRNU-M athymic nude mice (Taconic Farms), under anesthesia via ketamine/xylazine intraperitoneal injection, as described in detail [89]. To compare survival curves between mice implanted with CSCs treated with 0 Gy versus 4 Gy RT for each model, the required sample size was calculated using PASS Sample Size Software. The calculation was based on a one-tailed log-rank test with an alpha level of 0.10. With a sample size of 6 per group, there is 81% power to detect a difference in survival probabilities of 95% versus 5%. To account for possible losses, a total of 42 mice were used, n=7/group except for HF2303 for which n=6 mice were used for control and n=8 for irradiated CSCs. Following surgery to implant CSCs, mice were kept warm using a heating pad until they regained consciousness, returned to the cages and for 48h observed every 3h for any signs of distress, in which case buprenorphine can be administered. Subsequently, mice continued to be monitored daily and were weighted 3 times a week by a technician blinded to the experimental groups. Tumor growth was monitored by noninvasive bioluminescence imaging using IVIS Spectrum In Vivo Imaging System (Caliper Life Sciences) once a week. The primary endpoint of this study is symptom-free survival, and no mice were excluded from the study. Within 6 hours after the observation of any symptoms associated with tumor burden, which happened within 275 days post-implant, all 42 mice were euthanized by >5% isoflurane overdose, and after no signs of palpebral reflexes and breathing were observed, mice were decapitated as a second assurance of death. The symptoms included weight loss greater than 20% of body mass, cranial protrusion, sickness, weakening, inactivity, anorexia, hunched posture, sunken eyes, ruffled coat, abnormal head tilt, seizures, or ataxia.

#### Sublethal temozolomide and radiation treatment for total RNA and genomic DNA isolation

*Temozolomide*: GBM CSCs or SDCs growing in their respective media were treated in triplicate with the respective TMZ IC40 concentrations or control DMSO for 4 days. *Radiation*: Cells were treated in triplicate with ionizing radiation (4Gy) or mock radiation control (0Gy) and cultured for 5 days. For all treatment groups, cells were harvested, total RNA isolated using RNeasy Mini Kit (Qiagen) and used for RNA sequencing. Genomic DNA was isolated using QIAamp DNA kit (Qiagen) and used for DNA methylation profiling.

### RNA Sequencing

RNA quality assessment was performed using the Agilent RNA ScreenTape on the Agilent 2200 TapeStation. Library for RNA sequencing was prepared using TruSeq Stranded Total RNA LT (Illumina). 100 bps paired-end libraries were sequenced on the Illumina HiSeq 2500 following cBot clustering. Prior to alignment the sequencing adaptors were trimmed using FASTQ Toolkit v1.0 on Illumina’s BaseSpace portal. FastQC was used to generate quality metrics for assessment of FASTQ files [90]. RNA seq reads were aligned using TopHat v2.1.0 using default alignment parameters and GRCh38 reference genome [91, 92]. After alignment, the mapped reads were further filtered by mappability score (MAPQ ≥ 10) and sorted by genomic position of the reads using samtools-0.1.19 [93]. PCR duplicate reads were processed and removed using rmdup function (options -S) in samtools. The cleaned BAM files were further analyzed using Integrative Genome Viewer (http://software.broadinstitute.org/software/igv/), processed with R function “FeatureCount” to quantify reads on various RNA types from GencodeV28 annotation. Differential gene expression analysis was performed using the non-parametric algorithm “NOISeq” in R [94], with q = 0.8 and fold change >2 or <0.5 thresholds on TMM normalized data for treated vs. control comparisons, and q = 0.8 on RPKM normalized data for long term transcriptome changes in untreated SDCs vs CSCs. Gene set enrichment tests for differentially expressed gene lists were performed using Metascape 3.5, and adjusted p-value (q) < 0.05 [95].

### DNA methylation analysis

Genomic DNA was analyzed using Infinium Human Methylation 450K BeadChip system (Illumina), as described [86]. Raw methylation data was pre-processed for dye-bias normalization, detection p-value was calculated by comparing the signal intensity difference between array probes and a set of negative control probes on the array; probes with p-value greater than 0.01 filtered out. Beta values were calculated from pre-processed raw data using methylumi Bioconductor package in R [96]. Probes mapping to known SNPs and sex chromosomes were filtered out of dataset as recommended by Zhou et al. [97]. Beta values were corrected for batch effect using ComBat function within sva R package [98, 99]. Probes were aligned to hg38 and annotated using Illumina methylation 450k manifest [100]. Differentially methylated positions between experimental groups (n=3/group) were determined using champ.DMP function, and to determine differentially methylated regions we used the champ.DMR with Bumphunter method from the ChAMP package in ChAMPs R package [101] with raw p-values < 0.05, and adjusted p-value < 0.1. DMPs were annotated to genes using illuminaHg19 annotation employed by the ChAMPs package in R. Analyses to determine the intersect of DMPs, the genes mapped to DMPs and DEGs, within and between models, were carried out using GeneOverlap (version 1.34.0).

### Cellular oxygen consumption rate

Cellular oxygen consumption rate (OCR) was measured using a Seahorse XF Cell Mito Stress Test in a XFe24 Bioanalyzer (Agilent). Cells were plated at a concentration of 30,000 (HF2303) or 40,000 (HF2927) per well on poly-ornithine hydrobromide (Sigma # 4957) coated plates in NMGF media (CSC) or 2%FBS media (SDC) and treated for 4 days with DMSO or TMZ IC40 concentrations, prior to basal oxygen consumption measurements, performed according to the manufacturer’s instructions.

### Statistical analysis

Statistical analyses were performed using GraphPad Prism (v. 9.00) or R and are described in the results section for each experiment. Tumor growth in mouse PDX was evaluated by Kaplan-Meier survival curves, compared by the log-rank test with p<0.05 considered significant.

## Data availability

The data discussed in this publication have been deposited in NCBI’s Gene Expression Omnibus [102] and are accessible through GEO Series accession number GSE249637.

## Acknowledgments

We thank Kristyn Quenneville and Dr. Nouman Mughal for their assistance with reviewing and editing the manuscript.

## Author contributions

A.C.D. and D.R. conceived the study. K.B., Y.M., S.I., L.H., and A.B. performed cell culture, treatment response, and prepared cell lysate, DNA and RNA for profiling. C.M. and E.F.P. generated the RPPA data and performed initial analysis at Mason’s Center for Applied Proteomics and Molecular Medicine (CAPMM). AB performed RPPA data analysis. S.B. performed the radiation experiments. KG and DR performed the RNA sequencing and DNA Methylation at the Wayne State University Genomics Core. ID, LP and ON performed the RNAseq bioinformatics analysis. OD, TS and HN performed the DNA methylation bioinformatics analysis. N.P. and S.A. performed the OCR measurements. A.C.D., A.B. and O.N wrote the manuscript with input from co-authors.

## Supporting information

**S1_Fig. Alteration in glioblastoma cancer stem cell signaling in response to growth factors withdrawal and differentiation in 2% FBS.** A) Box plot representing changes in the levels of 66 proteins/PTM in response to 14-days culture in the absence of growth factors (NM), or upon serum differentiation (2%FBS), relative to CSC culture conditions (NMGF), using mean values from triplicate RPPA measurements for each of the 8 models. Plot represents n=507 data points, after proteins/PTMs which were not detected under all three culture conditions were filtered out for each model.

**S2_Fig. MGMT promoter methylation.** Heatmap representing β-values of 26 CpG sites mapping to MGMT promoter, for HF2303 and HF2927 CSC and SDCs in control, TMZ and RT treatment groups, in triplicate. Human astrocyte cell line was included as reference. The two probes widely employed in the clinic to determine MGMT promoter methylation status (“predictive probes”) indicate MGMT promoter hypermethylation in HF2927 and unmethylated status in HF2303 and human astrocytes cells. Differentiation status and treatment did not alter promoter methylation levels for either GBM line.

**S3_Fig. Alterations in cellular respiration in response to treatment.** A) Basal (B) and maximum (M) relative oxygen consumption rate (OCR) for CSCs and SDCs measured after a 4-day treatment with TMZ, or 4 days after treatment with one 4 Gy radiation dose. OCR values were normalized to CSC (left panel) and SDC (right panel) HF2303 basal levels (%). Measurements for n=2-3 repeats with means are shown. B) Reserve respiratory capacity was calculated as ratio of (mean maximum)/(mean basal) O2 consumption.

**S4_Fig. Short term treatment with TMZ and RT did not significantly alter DNA methylation pattern of glioblastoma CSCs and SDCs.** Principal component analysis for b-values for all treatment groups in triplicates for each cell line and differentiation status: HF2303 CSC (A), HF2303 SDCs (B), HF2927 CSCs (C), and HF2927 SDCs (D).

**S1 Table. Reverse Phase Protein Array (RPPA) Targets, Values, Signaling Pathways**

**S2 Table. Differentially expressed gene lists**

**S3 Table. Interferon signaling correlation group in glioblastoma (TCGA)**

**S4 Table. Differentially methylated probes**

