## Supplementary material for "Impact of genomic background and developmental state on signaling pathways and response to therapy in glioblastoma patient-derived cells": S1_Fig - S4_Fig

**SUPPLEMENTARY FIGURES**

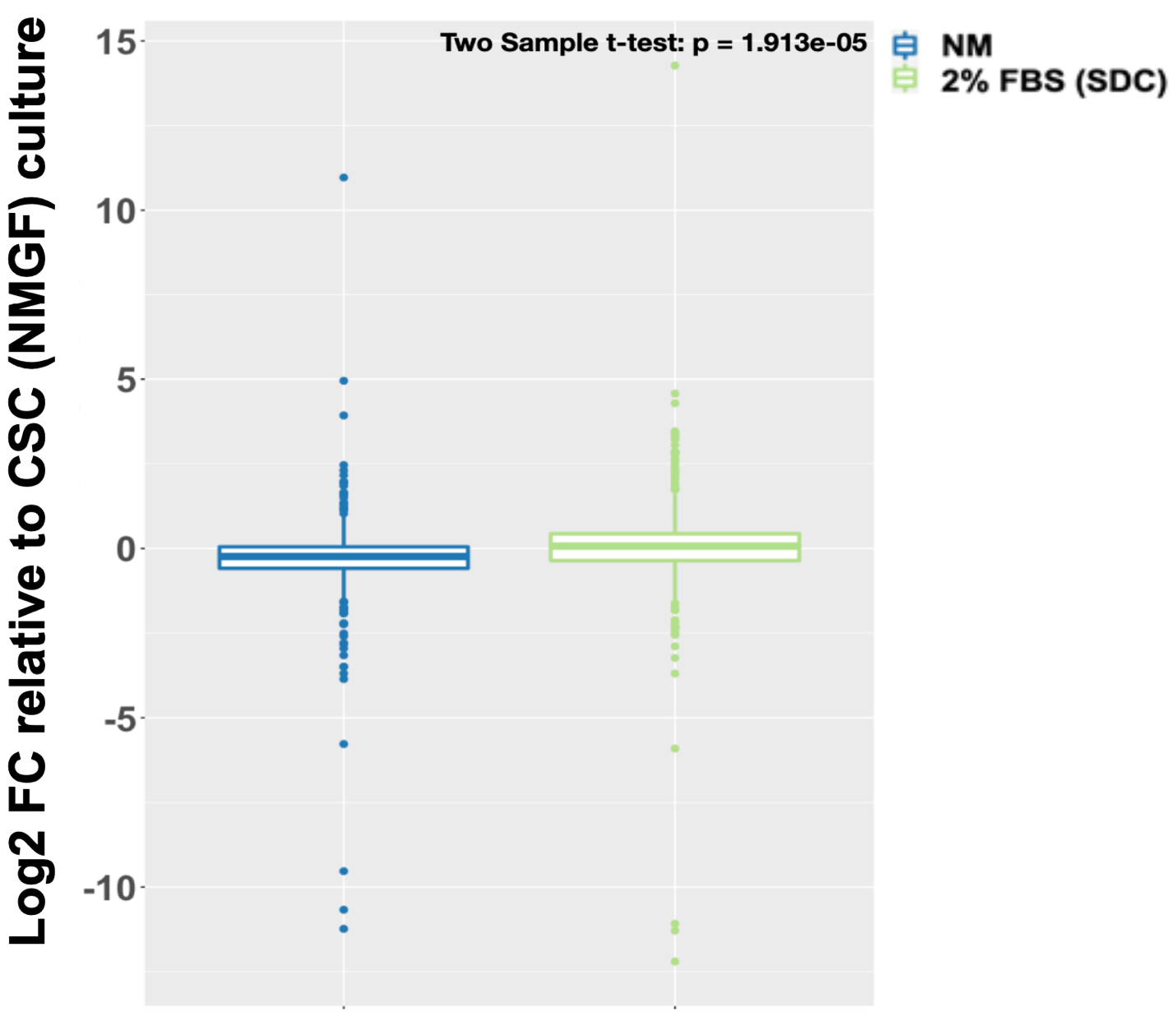

**S1_Fig. Alteration in glioblastoma cancer stem cell signaling in response to growth factors withdrawal and differentiation in 2% FBS.** A) Box plot representing changes in the levels of 66 proteins/PTM in response to 14-days culture in the absence of growth factors (NM), or upon serum differentiation (2%FBS), relative to CSC culture conditions (NMGF), using mean values from triplicate RPPA measurements for each of the 8 models. Plot represents n=507 data points, after proteins/PTMs which were not detected under all three culture conditions were filtered out for each model.

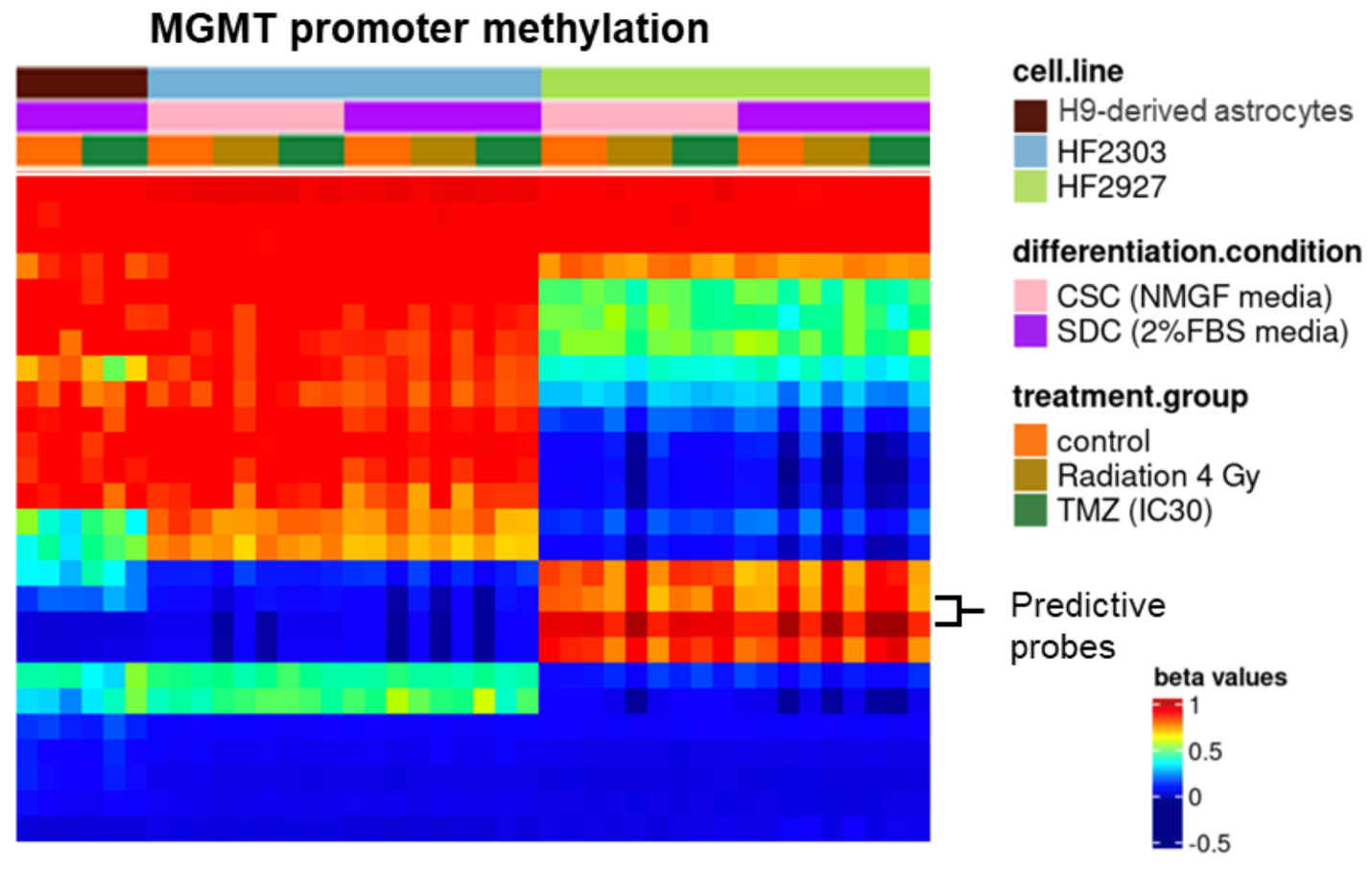

**S2_Fig. MGMT promoter methylation.** Heatmap representing β-values of 26 CpG sites mapping to MGMT promoter, for HF2303 and HF2927 CSC and SDCs in control, TMZ and RT treatment groups, in triplicate. Human astrocyte cell line was included as reference. The two probes widely employed in the clinic to determine MGMT promoter methylation status (“predictive probes”) indicate MGMT promoter hypermethylation in HF2927 and unmethylated status in HF2303 and human astrocytes cells. Differentiation status and treatment did not alter promoter methylation levels for either GBM line.

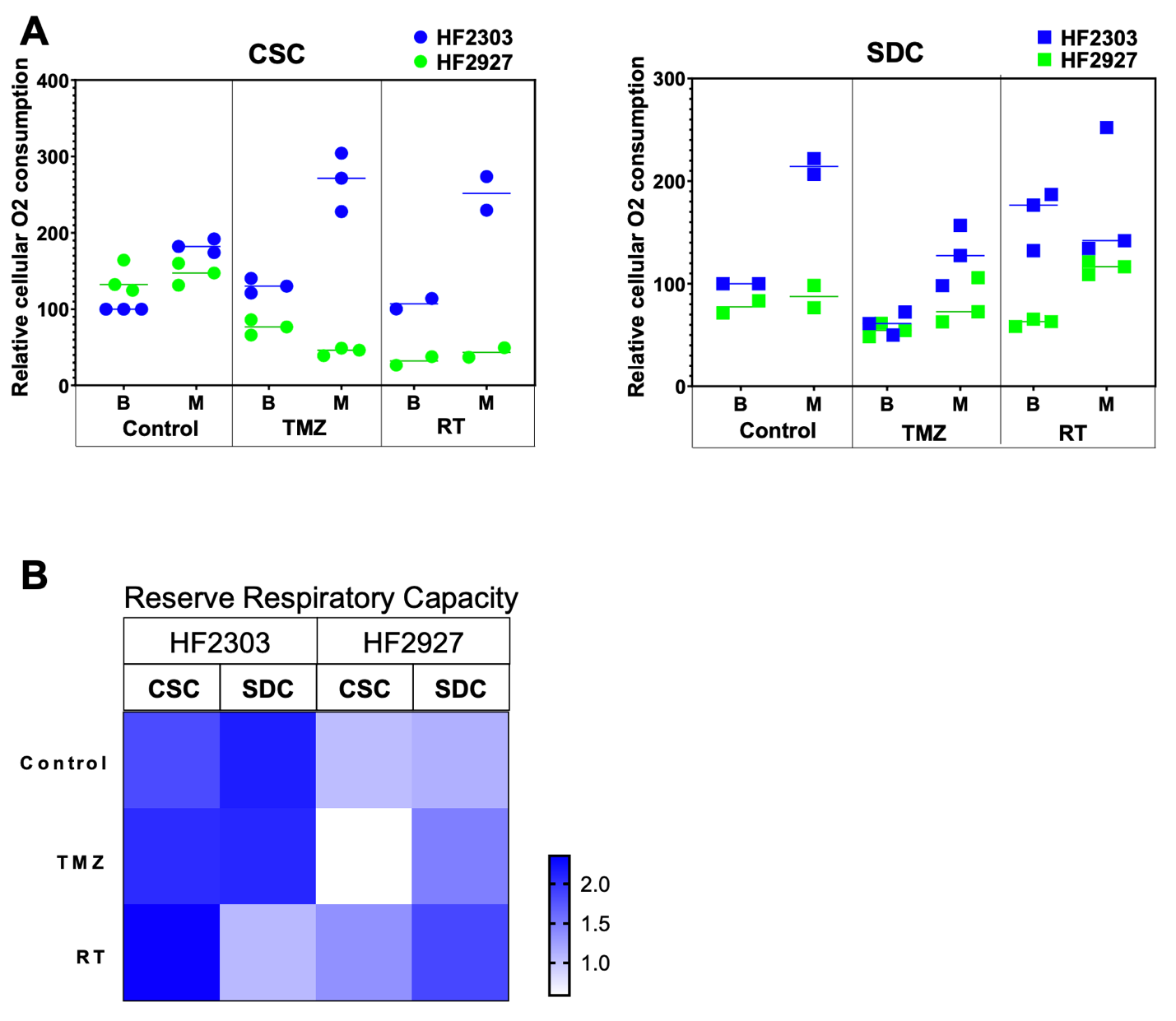

**S3_Fig. Alterations in cellular respiration in response to treatment.** A) Basal (B) and maximum (M) relative oxygen consumption rate (OCR) for CSCs and SDCs measured after a 4-day treatment with TMZ, or 4 days after treatment with one 4 Gy radiation dose. OCR values were normalized to CSC (left panel) and SDC (right panel) HF2303 basal levels (%). Measurements for n=2-3 repeats with means are shown. B) Reserve respiratory capacity was calculated as ratio of (mean maximum)/(mean basal) O2 consumption.

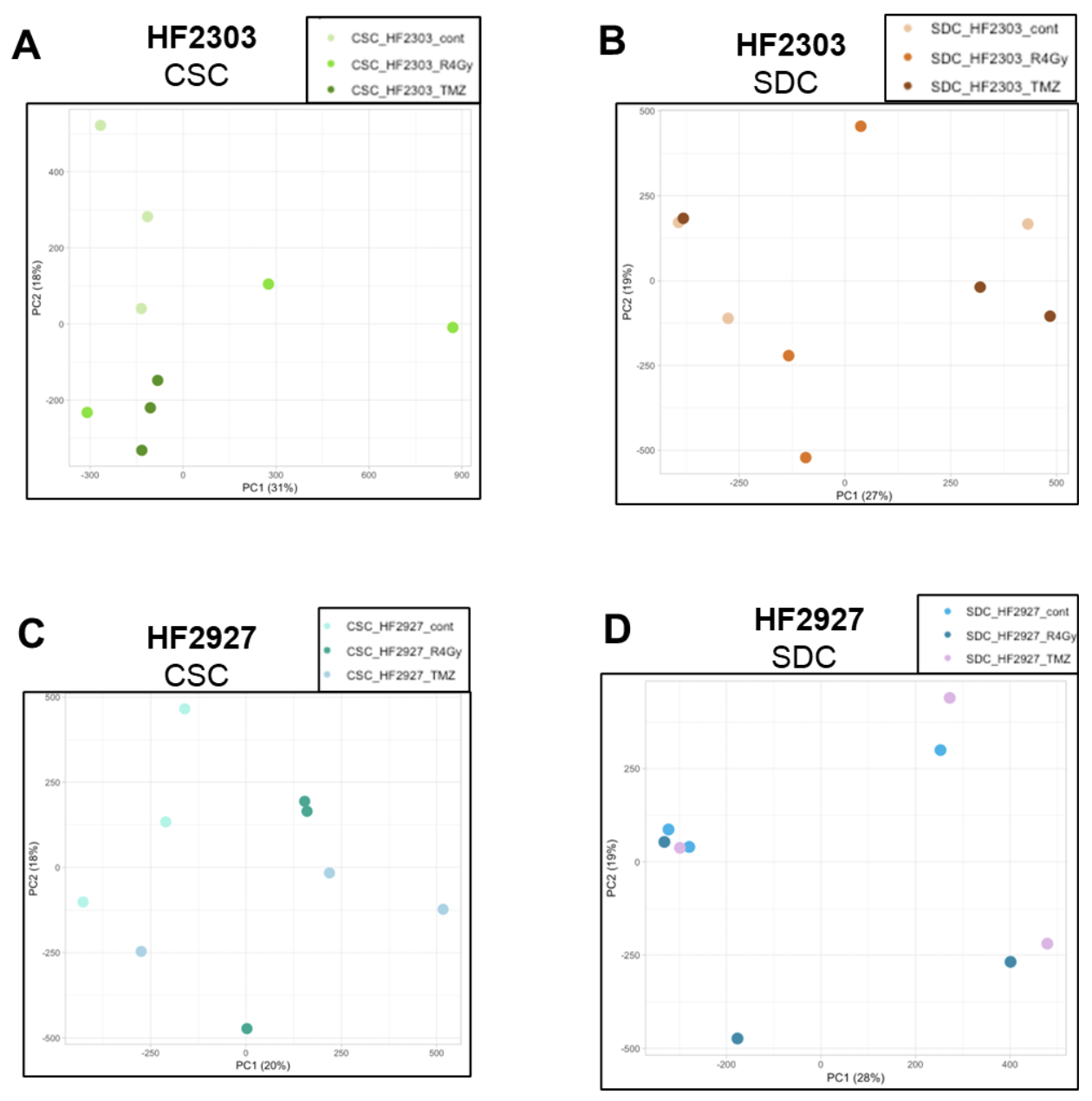

**S4_Fig. Short term treatment with TMZ and RT did not significantly alter DNA methylation pattern of glioblastoma CSCs and SDCs.** Principal component analysis for b-values for all treatment groups in triplicates for each cell line and differentiation status: HF2303 CSC (A), HF2303 SDCs (B), HF2927 CSCs (C), and HF2927 SDCs (D).
